# VARS-fUSI: Variable Sampling for Fast and Efficient Functional Ultrasound Imaging using Neural Operators

**DOI:** 10.1101/2025.04.16.649237

**Authors:** Bahareh Tolooshams, Lydia Lin, Thierri Callier, Jiayun Wang, Sanvi Pal, Aditi Chandrashekar, Claire Rabut, Zongyi Li, Chase Blagden, Sumner L. Norman, Kamyar Azizzadenesheli, Charles Liu, Mikhail G. Shapiro, Richard A. Andersen, Anima Anandkumar

**Affiliations:** Computing + Mathematical Sciences, California Institute of Technology; Division of Biology and Biological Engineering, California Institute of Technology; Division of Chemistry and Chemical Engineering, California Institute of Technology; Cherng Department of Medical Engineering, California Institute of Technology; Howard Hughes Medical Institute; USC Neurorestoration Center and the Departments of Neurosurgery and Neurology, University of Southern California; Rancho Los Amigos National Rehabilitation Center; NVIDIA; T & C Chen Brain-machine Interface Center, California Institute of Technology

## Abstract

Functional ultrasound imaging (fUSI) is a promising neuroimaging method that infers neural activity by detecting cerebral blood volume changes. It offers high sensitivity and spatial resolution relative to fMRI and is an epidural alternative to electrophysiology for medical and neuroscience applications, including brain-computer interfaces. However, current fUSI methods require hundreds of compounded images and ultrasound pulse emissions, leading to high computational costs, memory demands, and potential probe heating. We propose VARiable Sampling fUSI (VARS-fUSI), the first deep learning fUSI method to allow for different sampling durations and rates during training and inference by using neural operators. VARS-fUSI reconstructs high-quality fUSI images using 10 − 15% of the time or sampling rate needed per image while preserving decodable behavior-correlated signals. Additionally, VARS-fUSI offers efficient finetuning for generalization to new animals and humans. Demonstrated across mouse, monkey, and human data, VARS-fUSI achieves state-of-the-art performance, enhancing imaging efficiency by significantly reducing storage and processing needs.

## Introduction

Functional ultrasound imaging (fUSI)^1,2^ is a promising and minimally-invasive neuroimaging technique that hemodynamically measures neural activity from outside of the dura, unlike intracortical implants. Similar to fMRI, fUSI infers population-level neural activity from changes in cerebral blood volume (CBV) by leveraging the concept of neurovascular coupling. However, fUSI has a higher sensitivity (1 mm/s velocity) and spatial resolution (100-300 *µm*) than fMRI. Additionally, it can epidurally image from a large field of view (≥ 12.8 mm × 16 mm, *W* × *H*), providing a more minimally invasive alternative to electrophysiology. fUSI’s low cost and portability, given the small size of ultrasound probes, also allow for bedside use ^3,4^ and simultaneous use in freely moving animals^5–7^ and human subjects^8^. These advantages make fUSI an optimal tool for minimally invasive examination of brain activities in the clinic and during neuroscience studies. Clinically, fUSI has already been used to map brain activity and vasculature intraoperatively in humans for tumor removal^9^ and monitor functional connectivity and epileptic activity in human neonates at bedside^3,4^. Several prior studies have used fUSI to record neural activity in behavioral tasks in freely moving rodents^5^, non-human primates^6,7^, and humans^8^. Moreover, fUSI has been increasingly used for brain-computer interfaces (BCIs)^6–8^, a technology that decodes and converts brain signals into computer commands, demonstrating that fUSI has potential to power the next generation of minimally invasive BCIs.

Despite its promise, real-time computational demands limit the adoption of fUSI. To produce a single power Doppler (DOP) image, hundreds of ultrasonic planewave emissions are transmitted and received at ultrafast rates, capturing backscattered signals from red blood cells, tissue, and noise. These signals are beamformed and coherently compounded to produce intermediate in-phase and quadrature (IQ) frames. Hundreds of compounded frames are processed using singular value decomposition (SVD)^10–13^ to remove tissue motion and isolate blood volume signals. The filtered frames are then squared and averaged to generate a DOP image, which reflects changes in CBV. This pipeline requires high pulse emissions and large storage, which limits imaging framerates and can cause probe heating exceeding FDA safety guidelines, ultimately restricting the capabilities and broader application of fUSI.

Two primary computational obstacles contribute to these constraints: a) computational complexity and memory storage challenges due to the number of intermediate IQ frames that must first be acquired and beamformed per DOP and b) SVD, used for tissue motion filtering, involves slow and expensive computations, does not scale for large datasets, and requires domain experts to tune the filtering parameters. Deep-learning-based approaches such as Deep-fUS ^14^ have aimed to address these challenges with their scalable graphical processing unit (GPU) infrastructures. However, they operate on a fixed framerate, and their applicability remains limited in practice when a change or reduction of the framerate is needed.

### Our approach

We propose VARiable Sampling Functional Ultrasound Imaging (VARS-fUSI)^1^. VARS-fUSI is a deep-learning-based framework that improves the efficiency of fUSI by reducing the number of compounded frames needed to reconstruct DOP images. This allows users to either a) increase the fUSI imaging framerate or b) reduce the data sampling rate needed per DOP. This alleviates the computational, memory, and communication complexity of fUSI technology.

VARS-fUSI is based on neural operators, which map between infinite-dimensional functions^15–17^. Neural operators are applied in solving partial differential equations (PDEs)and inverse problems^18^, with diverse scientific and engineering applications^19^, including MRI^20^, lung ultrasound^21^, computational fluid dynamics^22^, net-zero climate^23^, seismology^24^, and weather forecasting^25^. For MRI, neural operators have recently enabled the creation of a unified model applicable to varying frequency sampling patterns^20^. For lung ultrasound, they have facilitated bypassing a beamforming step by analyzing directly the radio-frequency signals^21^.

A notable property of neural operators is their ability to process varying discretizations, enabling flexible adjustments in hardware sampling rates or acquisition durations without compromising imaging quality. VARS-fUSI is the first fUSI method based on neural operators to fully leverage this capability. Unlike traditional deep learning approaches, it treats complex IQ signals as continuous functions in time, allowing VARS-fUSI to process data with sampling rates or durations different from the training set, without retraining. VARS-fUSI also advances beyond prior neural operator applications in MRI^20^ and lung ultrasound^21^ by implementing decomposed temporal and spatial processing in each deep layer. This enables flexible changes in sampling time and rate. VARS-fUSI includes a spatial guidance module that estimates blood flow locations using SVD-based spatiotemporal filtering. Combined with temporal filtering, this design enables a generalizable mapping that is agnostic to acquisition framerate.

We applied VARS-fUSI to fUSI brain images collected from mouse, monkey, and human and demonstrate VARS-fUSI state-of-the-art performance in achieving high-quality reconstructions of fUSI data using less temporal data. We showcase VARS-fUSI’s superior generalization to unseen experiment sessions in new animals and across species while being trained on image frames from only mice. Additionally, we offer a fast and efficient finetuning procedure to further improve its generalization to new animals; this can be achieved on a single GPU with less than 24 GB of memory in the order of minutes, making VARS-fUSI easy to implement, adapt, and apply for users. Finally, we show that VARS-fUSI can preserve spatiotemporal features necessary for behavioral decoding in monkeys and humans when reducing the amount of time (reduced-time) or the sampling rate (reduced-sampling) needed to produce one DOP image. This demonstrates VARS-fUSI’s potential for application in BCIs.

For the reduced-time approach, which takes a limited number of consecutive frames from the beginning of each DOP, the Doppler imaging framerate can be increased by a factor of ×10. This provides more image samples per unit of time and can be used to improve VARS-fUSI-powered BCIs. Reduced-sampling VARS-fUSI increased the sampling efficiency of imaging by reducing the IQ frame sampling rate by 90%. Thus, beyond VARS-fUSI’s ability to construct DOP images from less data, VARS-fUSI is the first deep learning framework of its kind that can operate at a sampling rate or duration different from the trained setting due to its temporal operator learning feature, having broad potential in improving fUSI technology and its applications.

Overall, VARS-fUSI provides: a) a training pipeline for experimental datasets (fig. 1a), b) an inference framework for accelerated online imaging capable of varying data acquisition time or sampling rate (fig. 1b), c) a fast fine-tuning procedure for improved session-specific precision (fig. 1c), and d) a low-latency and efficient imaging tool that preserves decoding information for BCI-based applications in monkeys and humans (fig. 1d).

**Figure 1:**
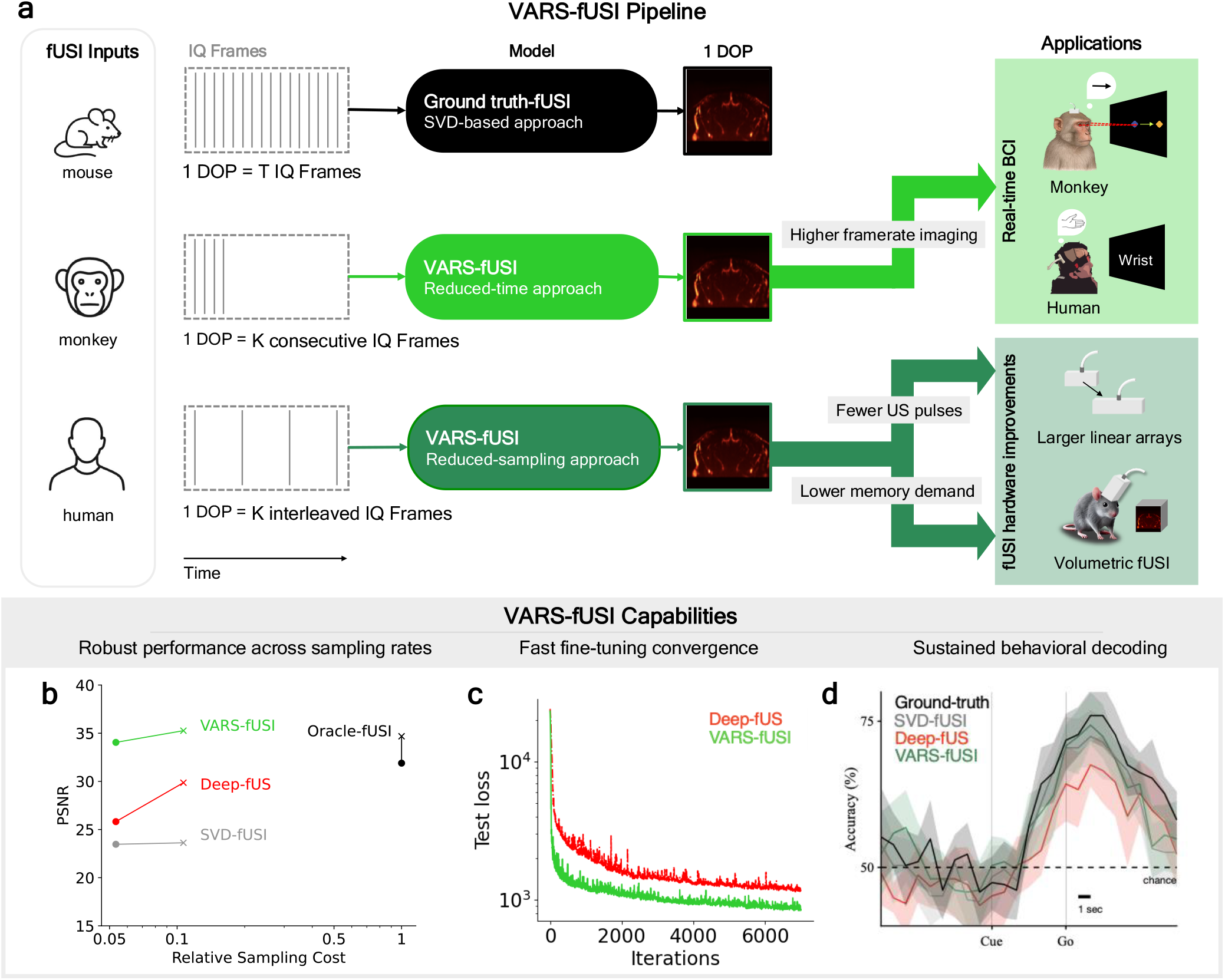
VARiable Sampling Functional Ultrasound Imaging (VARS-fUSI). **a**, VARS-fUSI pipeline. The ground-truth fUSI approach uses SVD-based filtering to construct 1 DOP image from *T* beam-formed IQ complex signals. VARS-fUSI reduces the number of input IQ frames from *T* to *K* ≪ *T*, and outputs a DOP image, which is the average power of the filtered and decluttered IQ frames. VARS-fUS takes the first *K* consecutive frames at the original sampling rate (reduced-time approach), or takes *K* frames within the same time period *T* as one ground-truth DOP at a lower IQ sampling rate (reduced-sampling approach). VARS-fUSI is applied to mice, monkeys, and humans (for VARS-fUSI’s architecture, see fig. S1). VARS-fUSI is trained by computing a loss based on the similarity between the output and the desired DOP image, outputs by the ground-truth fUSI method. The reduced-time VARS-fUSI enables increased framerate imaging; this approach does not change the discrete fUSI sampling rate but rather generates one DOP from smaller chunks of IQ frames in time. Reduced-time VARS-fUSI has promise in offering low-latency real-time BCI across different species. Reduced-sampling VARS-fUSI reduces the amount of data needed per DOP and allows for reduced ultrasonic emissions and data storage requirements. This has the potential to make fUSI technology more efficient as data requirements increase such as for larger linear arrays and volumetric fUSI. **b**, Compared to the Ground-truth and Oracle-fUSI, VARS-fUSI reduced the sampling cost by 90%. Moreover, VARS-fUSI can take a varying number of IQ frames as input and shows robust performance when reducing the sampling rate from the original trained rate (×) to its half-rate (○). Deep-fUS^14^ does not support a change in the number of frames, and its performance decays when operating at a reduced framerate compared to the original trained rate (see fig. 3). **c**, VARS-fUSI has accelerated fine-tuning compared to Deep-fUS. **d** While reduced-sampling VARS-fUSI improves the sampling efficiency by 90%, it preserves behavioral information for decoding in BCI; VARS-fUSI shows a more reliable and consistent decoding performance across sessions than Deep-fUS (see figs. 4 and 5).

## Results

### Deep neural operator framework for accelerated and efficient fUSI

VARS-fUSI (figs. 1 and S1) is a deep learning framework that offers more efficient fUSI from reduced compounded frames. Currently employed methods require hundreds of frames (e.g., 250 or 300) to achieve highquality fUSI images. VARS-fUSI maintained this performance using only 10 − 15% of the number of frames (e.g., 32 frames). We considered two approaches to achieve this sampling efficiency: a) reduced-time, taking the first *K* limited frames to construct the image, and b) reduced-sampling, taking frames within the same *T* -frame duration but at a lower IQ sampling rate. VARS-fUSI was GPU-enabled with parallelized processing over multiple images.

VARS-fUSI consists of three phases: training, fine-tuning, and inference. Given the *K* IQ frames input, VARS-fUSI was trained to minimize the error between VARS-fUSI’s output and the ground-truth DOP image using loss computation and backpropagation. Training was conducted on multiple mouse recording sessions, incorporating deep learning techniques such as data augmentation for improved performance. We publicly release our trained model for application to new imaging sessions and animals, with an optional fine-tuning stage. Fine-tuning required only a few minutes of data at the start of an experiment but was not always necessary, as we demonstrate that VARS-fUSI generalized across sessions without fine-tuning. During inference, VARS-fUSI reconstructed fUSI images from limited IQ frames in a fraction of a second.

VARS-fUSI integrates deep learning and operator learning (fig. S1) using a U-shaped architecture^26,27^ that applies decomposed spatial and temporal filtering to the beamformed complex IQ signal. VARS-fUSI learns tissue motion and blood flow statistics through this decomposition, a crucial aspect often overlooked in deep learning approaches. Spatial filtering is achieved via convolutional blocks, while temporal filtering is achieved globally using the Fourier neural operator (FNO)^16^, making the model discretization-agnostic. Three distinct characteristics enhance VARS-fUSI’s generalizability. First, unlike prior deep learning methods^14^, it preserves the complex nature of the IQ input signal. Second, it captures global frequency-based features essential for filtering slow-moving tissue components from blood flow, allowing it to operate at different sampling rates than those used during training. This improves memory and communication efficiency in fUSI hardware. Third, VARS-fUSI is spatially guided on the blood flow region by an SVD-fUSI module^13^.

We used SVD-processed DOP images, a widely used tissue decluttering method, as the ground truth. These ground-truth images were generated by applying SVD to the full set of *T* frames, removing tissue clutter across all time points, and averaging power over the entire set. As SVD filtering is the gold standard in fUSI for neuroscience, our goal was to match its image quality while using fewer IQ frames. The central frequency of fUSI transmissions varied across species to balance depth and resolution, and the total number of *T* frames was species-dependent, based on optimized parameters from prior studies^6–8^. To assess image quality under the *K*-frame constraint imposed by VARS-fUSI, we introduced two baselines: Oracle-fUSI and SVD-fUSI. Oracle-fUSI averages only the filtered *K* frames but uses all *T* frames for SVD-based tissue removal, serving as an upperbound reference. However, it is impractical as it assumes full-frame access. In contrast, SVD-fUSI is a practical baseline, using only the limited *K* frames for both clutter removal and averaging. By comparing VARS-fUSI to both baselines, we assessed its ability to reconstruct high-quality DOP images from significantly fewer IQ frames.

### VARS-fUSI is the state-of-the-art (SOTA) fUSI that runs efficiently at a faster framerate

We demonstrated the imaging quality of VARS-fUSI using the reduced-time approach. We trained VARS-fUSI on a fUSI imaging dataset (MultiMice) from five mouse brains acquired through intact skull and skin, where the target image to construct was the DOP image provided by SVD-based fUSI at the original framerate. We refer the reader to table S2 for more information on the data and to Methods for runtime analysis.

#### Model comparisons

The ground-truth images were constructed using *T* = 300 IQ frames, while VARS-fUSI operated with only the first *K* = 32 frames. In addition to SVD-fUSI, we compared VARS-fUSI with deep-learning-based methods: DnCNN^28^, Deep-fUS-base, and Deep-fUS ^14^ (fig. 2). DnCNN, a convolutional neural network originally designed for image denoising, processes only the real component of IQ frames^28^. Deep-fUS-base, an initial, basic version of Deep-fUS, lacks 3D convolution over time and space at the beginning of its architecture^14^. Deep-fUS treats IQ frames as channels, ignoring temporal aspects, and includes a single 3D convolutional block at the starting layer to mix frames temporally. Deep-fUS is currently the best-published deep learning method available for reconstructing fUSI data from sparse data. All models were trained similarly to VARS-fUSI, ensuring that any performance differences reflect architectural advantages. We evaluated performance using peak signal-to-noise ratio (PSNR, ↑), structural similarity index measure (SSIM, ↑), and normalized mean square error (NMSE, ↓) (see Evaluation Metrics in Methods). Using the MultiMice test set, VARS-fUSI outperformed competing models, achieving SOTA results (fig. 2a,b).

#### Importance of temporal filtering

First, we showed that reducing the number of IQ frames makes it harder to distinguish tissue clutter from blood flow, leading to poor SVD-fUSI performance. VARS-fUSI, trained to capture temporal correlations from limited IQ frames, improved upon SVD-fUSI and closely matched Oracle-fUSI. Second, effective data-driven filtering must leverage the inherent differences between blood flow signals and tissue clutter to ensure generalization. We compared VARS-fUSI to DnCNN^28^ and Deep-fUS-base^14^, and showed that VARS-fUSI outperformed both (fig. 2b). DnCNN and Deep-fUS-base lack temporal processing, limiting their ability to generalize across multiple brains (fig. 2a, Deep-fUS-base’s performance was similar to DnCNN). These findings highlight the crucial role of temporal filtering in achieving a generalizable fUSI framework.

**Figure 2:**
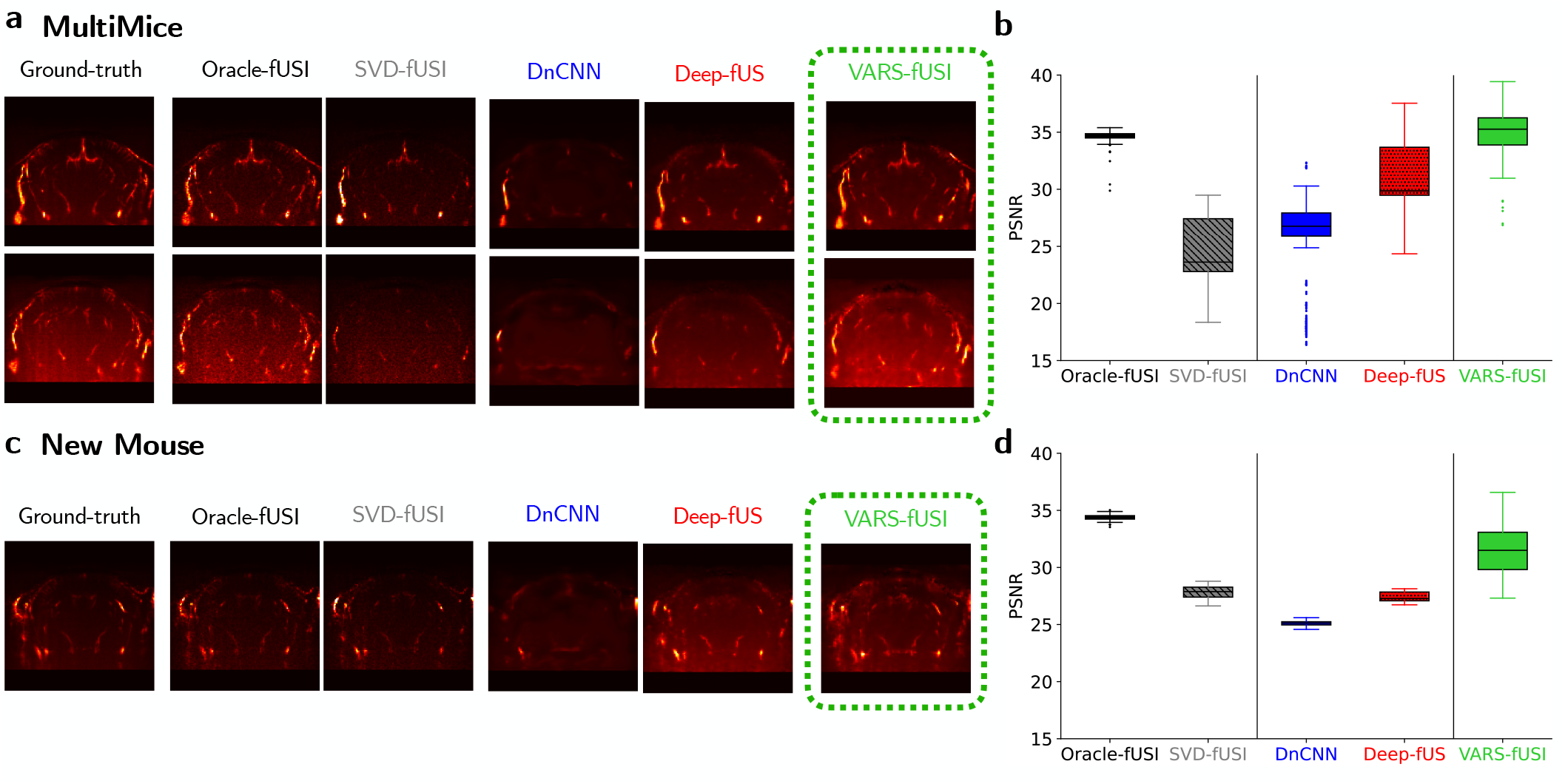
Outperformance generalization of VARS-fUSI on unseen frames from MultiMice and new mouse (Mouse X) for the reduced-time scenario. **a**, Qualitative analysis on the test set (unseen frames) from the MultiMice dataset. An example from two distinct mouse is shown. From left to right, columns represent: Ground-truth with *T* = 300 frames, Oracle-fUSI, SVD-fUSI^10,11,13^, DnCNN^28^, and Deep-fUS^14^, and VARS-fUSI for the reduced-time approach with *T* = 32. **b**, Quantitative comparison of VARS-fUSI with baselines in terms of PSNR on the test set, MultiMice (see also fig. S2a). **c**, Qualitative example of the performance on Mouse X for activity during visual stimulus presentation. **d**, Quantitative results on the new mouse (Mouse X) session M2 (see also fig. S2b,c).

#### Comparison to the best available method

Compared to Deep-fUS^14^, VARS-fUSI showed superior performance (fig. 2a,b; see also fig. S2). VARS-fUSI captured fine spatial details often missed by fully data-driven methods such as Deep-fUS. Our results highlight the importance of three key features unique to VARS-fUSI: 1) processing complex IQ frames rather than only the real component, 2) incorporating temporal processing layers, and 3) integrating spatial guidance and filtering in deep layers (see fig. S3a). Overall, unlike Deep-fUS, VARS-fUSI maintained strong generalization performance even in limited data settings (fig. S3b).

#### Generalization to a new mouse without fine-tuning

Next, we evaluated the generalization of VARS-fUSI for imaging on a new mouse, Mouse X, whose brain images were not included during training. We analyzed one experimental session containing DOP images from resting state (Mouse X M1) and during visual stimulation (Mouse X M2). In M2, a stimulus was applied every 10 DOP images, using 300 IQ frames per DOP (fig. 2c,d). VARS-fUSI outperformed SVD-fUSI in NMSE and PSNR. Deep-fUS failed to capture detailed blood flow information, performing worse than VARS-fUSI.

We attribute VARS-fUSI’s superior generalization to two key factors: 1) unlike Deep-fUS, it processes complex IQ signals, and 2) it incorporates spatial guidance via the SVD-fUSI module, which Deep-fUS lacks. To validate this, we conducted additional experiments using only the real component of the IQ input (fig. S4). Since spatial guidance requires complex data, it was omitted in this setting. First, VARS-fUSI performance declined without the spatial guidance module. While initial results on the MultiMice test set suggested that VARS-fUSI-real without guidance (w/o G) slightly outperformed VARS-fUSI w/o G, a closer analysis showed that the complex-input version generalized significantly better (fig. S4b,c), emphasizing the value of retaining the imaginary component.

### Variable sampling and duration with VARS-fUSI

**D**espite VARS-fUSI’s SOTA performance, a major challenge limiting the widespread adoption of deep learning for fUSI in practice is the difficulty of processing data with sampling rates or durations that differ from the original trained setting without re-designing or re-training a new model. This section highlights VARS-fUSI’s ability to process varying numbers of input time frames.

#### Variable rate generalization

An important advantage of VARS-fUSI over deep neural networks is its use of neural operators for global temporal processing rather than channel-wise processing of temporal frames. This enables VARS-fUSI to operate at 0.5 × (or [0.5 − 1) ×) the trained IQ sampling rate, skipping every alternate frame, resulting in an additional 50% reduction in computational, memory, and communication costs without performance degradation. We tested this on the MultiMice test set for the reduced-time approach. We inputted IQ frames at half the trained framerate, leading to a decrease in the number of time frames from 32 (1×) to 16 (0.5×). While there was a slight performance drop in Oracle-fUSI, SVD-fUSI, and VARS-fUSI due to the availability of fewer frames, DnCNN and Deep-fUS experienced a much sharper performance decline (fig. 3a,b).

**Figure 3:**
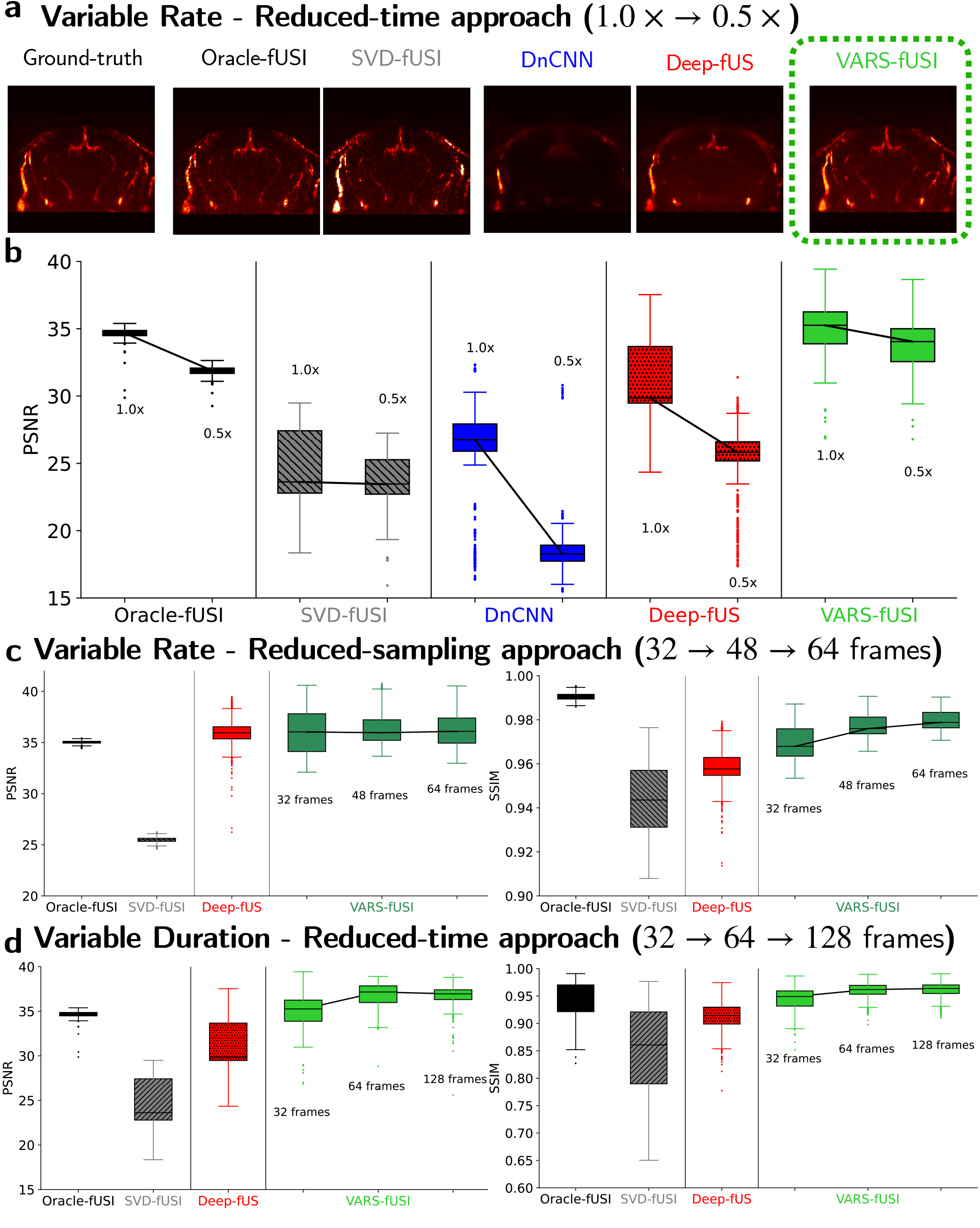
Demonstration of VARS-fUSI capabilities in changing sampling rate or duration. **a-b**, Changing the sampling rate from the original trained rate (1×) to half rate (0.5×) for the reduced time scenario when *K* = 32. Ground-truth DOP is computed using SVD de-cluttering based on 300 IQ frames. For 0.5× scenario, the input frames are interpolated from 16 to 32 frames. This is to be able to run methods not supporting input of different sizes. **c**, Varying the sampling rate for reduced sampling approach to acquire 32, 48, or 64 frames. Oracle-fUSI, SVD-fUSI, and Deep-fUS are shown at the original rate with *K* = 32 frames. Deep-fUS cannot change the number of input frames. The models are fine-tuned on the first 120 images of *S*1 from the new Monkey and tested on the rest of the hours-long session. **d**, Varying the duration of data acquisition at a fixed sampling rate for the reduce time approach. The number of frames is increased from the original trained *K* = 32 to 48 and 64. Oracle-fUSI, SVD-fUSI, and Deep-fUS are shown at the original duration with *K* = 32 frames. Deep-fUS cannot change the number of input frames. The models are tested on the MultiMice dataset.

We then considered the reduced-sampling approach and showed that the sampling rate could be increased for improved performance. In this case, DnCNN and Deep-fUS could not process a different sampling rate from the original training set, and thus were not applicable. VARS-fUSI was capable of increasing the rate from 32 frames to 48 and 64 frames. This resulted in improved imaging quality in performance metrics, though they were below the Oracle-fUSI (fig. 3c).

#### Variable duration generalization

VARS-fUSI can also be applied to inputs of different durations at a fixed sampling rate. Given the original 32 frames shown for Oracle-fUSI, SVD-fUSI, and Deep-fUS, we increased the duration of data inputted to VARS-fUSI to 64 and 128 frames. This led to improved performance (fig. 3d). Similar to the variable sampling scenario, Deep-fUS lacks the capability of accepting IQ frames of different temporal lengths, hence, it was shown only in the original trained 32 frames.

### Behavioral decoding using VARS-fUSI and practical generalization on new sessions and animals

A major limitation of deep learning methods is poor generalization beyond the training distribution, known as domain shift^29^. In medical imaging, this challenge, termed out-of-distribution (OOD) generalization, is particularly evident across sessions or subjects^30^. To address this, we extended VARS-fUSI’s reliability and usability across species and sessions through an efficient fine-tuning process. Starting from a VARS-fUSI model trained on MultiMice, we fine-tuned it for the target session or species. We demonstrate that this can be done quickly on a single GPU, enabling application to imaging from new mice, monkeys, and humans. Beyond fine-tuned sessions, we also show that VARS-fUSI generalizes to unseen sessions from the same subject and retains behavior-correlated information, supporting its use in BCI applications.

fUSI-based BCIs are an emerging technology capable of decoding movement and planning signals, enabling control of robotic limbs and assistive devices more minimally invasively than traditional electrophysiological BCIs^6–8^. We propose using VARS-fUSI to power BCIs by a) increasing imaging framerate through the reduced-time approach and b) improving sampling efficiency via reduced-sampling VARS-fUSI. Here we show that, unlike prior deep learning methods, VARS-fUSI can produce high-quality DOP reconstructions and preserve behavior-related information crucial for decoding complex behaviors in BCIs in both monkey and human experiments.

#### Monkey experiment imaging

We applied VARS-fUSI to functional ultrasound neuroimaging in Rhesus macaques (fig. 4), analyzing three sessions—S1, S2, and S3. VARS-fUSI, pre-trained on MultiMice, was fine-tuned using 120 DOP images from the start of S1. We then evaluated its performance on the remaining frames of S1 (fig. 4b,c) and the full sessions of S2 and S3 (figs. S5 and S6), which were not seen during fine-tuning. The ground-truth DOP images were computed using *T* = 250 IQ frames, while VARS-fUSI, SVD-fUSI, and Deep-fUS used only *K* = 32 frames. SVD-fUSI, while capturing general blood flow, lost fine details due to thresholding sensitivity with limited frames, resulting in high NMSE variance and low PSNR. VARS-fUSI outperformed both SVD-fUSI and Deep-fUS, showing lower variance and image quality close to Oracle-fUSI, with reduced noise in unseen sessions. VARS-fUSI’s generalization also held in the reduced-sampling setting for S1 (fig. 4f,g), and similarly for imaging and decoding in S2 and S3 (fig. S5a,b and fig. S6c,d).

**Figure 4:**
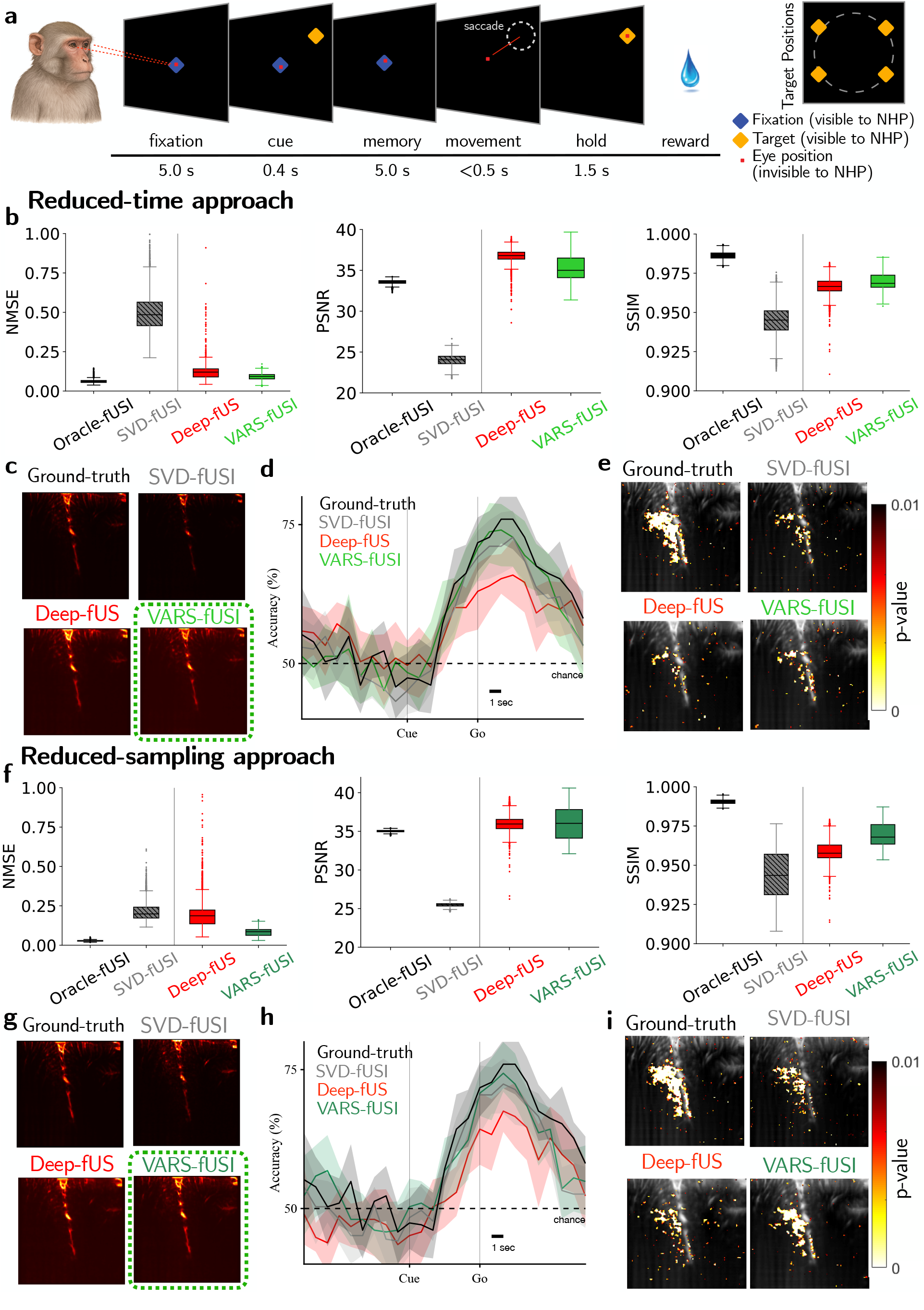
VARS-fUSI-powered BCI to decode thoughts from monkey. Decoding is performed using a 4 s sliding window. **a**, Memory guided saccade task in session S1 for decoding varied directions for target/fixation points. ±1000 ms of jitter for fixation and memory periods; ±500 ms of jitter for hold period. The red square signifies the monkey’s line of sight. **b**, Quantitative image similarity metrics for the reduced-time approach. **c**, Qualitative example images of each models’ performance using the reduced-time approach. **d**, Horizontal direction decoding performance for S1 when models produce DOP images via reduced-time approach (i.e., 8 DOP images at 32 IQ framerate from 250 IQ frames). The ground-truth constructed only one DOP image per 250 IQ frames (per second). The vertical lines denote the Cue and Go onsets. **e**, Statistically significant task-correlated activation maps with *p <* 0.01 of each method using the reduced-time approach calculated by performing voxelwise t-tests. **f**, Quantitative image similarity metrics for the reduced-sampling approach. **g**, Qualitative example images of each models’ performance using the reduced-sampling approach. **h**, Horizontal direction decoding performance from S1 for reduced-sampling approach. Models constructed one DOP image using 32 out of 250 IQ frames sampled at 7× less rate compared to the original framerate. **i**, Statistically significant task-correlated activation maps with *p <* 0.01 of each method using the reduced-sampling approach calculated by performing voxelwise t-tests.

#### Monkey experiment decoding

We recorded fUSI data from the lateral intraparietal sulcus (LIP, a cortical area critical for eye movement planning^31–33^) of a rhesus macaque performing two eye saccade tasks: one that varied saccade direction from a fixed starting point (S1 and S2,fig. 4a), and another that varied both starting location and saccade amplitude (S3,fig. S6a). The goal was to decode movement planning prior to action. We used an offline decoding pipeline with dimensionality reduction and linear decoding to predict planned movement direction and distance after complete data collection (see Methods).

Reduced-time VARS-fUSI increased the DOP imaging rate by ≈ 8 ×, generating 8 DOP images using 32 IQ frames in the 1 second typically needed to produce just one. This higher framerate is valuable for implementing online decoders, highlighting the potential for real-time applications. Reduced-sampling VARS-fUSI produced one DOP image at the ground truth rate but with 7× lower IQ sampling, making it ideal for large fields of view and volumetric imaging, where memory and ultrasound emissions are limiting factors.

To assess VARS-fUSI’s ability to retain decodeable behavioral information, we compared the decoding performance and task-activation statistical maps of ground-truth, SVD-fUSI, Deep-fUS, and VARS-fUSI. We evaluated decoding performance over a 4-second sliding window. Our pipeline also provides DOP augmentations, which imitate movement during imaging, prior to decoding to further improve VARS-fUSI performance (fig. S7).

In the reduced-time scenario, we averaged eight images per second to match the four DOP images per window used in the ground truth during decoding. We found that VARS-fUSI performed similarly to ground truth and SVD-fUSI while Deep-fUS had worse performance (fig. 4d; for other sessions, see fig. S6b). fig. 4e shows the task-related activation maps, confirming that VARS-fUSI maintains spatial representations of behavioral information in the brain.

In the reduced-sampling approach, all methods produced four DOP images per window. For S1, using the same window size as in the reduced-time scenario, VARS-fUSI maintained minimal decline in performance despite achieving an 87% more efficient sampling rate, while Deep-fUS and SVD-fUSI showed a slight decline in decoding (fig. 4h; for other sessions, see fig. S6c). Deep-fUS’ lower accuracy was likely due to the model’s high variance, failing to consistently generate highquality DOP images. Additionally, VARS-fUSI statistical significance maps (fig. 4i) were closest to the ground-truth in comparison to SVD-fUSI and Deep-fUS.

#### Human experiment imaging

Next, we demonstrated VARS-fUSI’s ability to generalize to imaging from a human subject with an acoustic cranial implant for the reduced-time approach (fig. 5). In these experiments, we used the VARS-fUSI that was already trained on MultiMice and finetuned it only on the first few images of a human session. To evaluate imaging quality, the desired ground-truth DOP images were computed using *T* = 300 IQ frames. We focused on performance in two separate human sessions (H1, H2).

**Figure 5:**
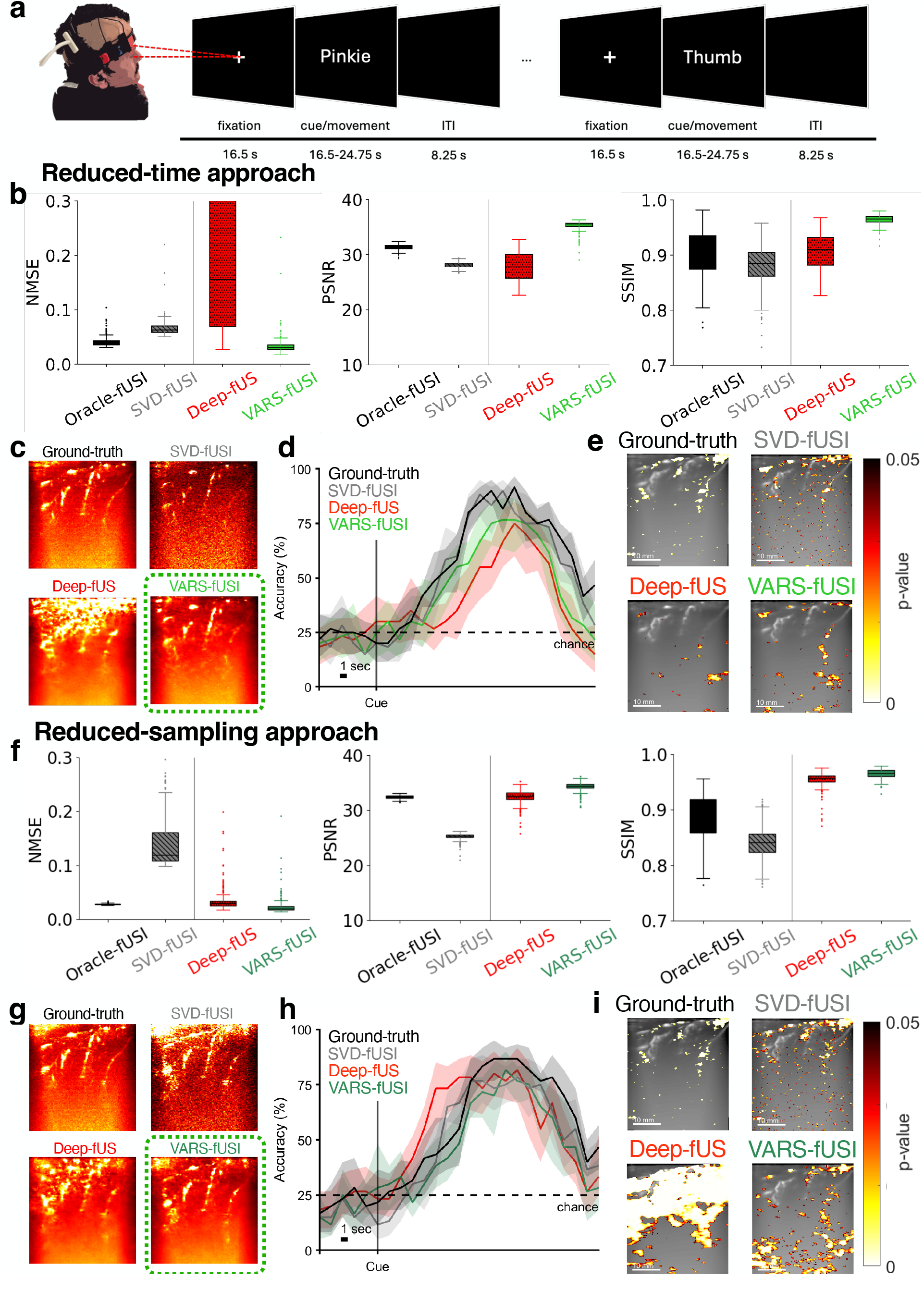
VARS-fUSI-powered BCI to decode planning from human. **a**, Instructed delay motor task for human experiments. The participant was asked to fixate on a central fixation cross and then perform the cued movement upon cue. The participant was asked to repeat the movement until the cue disappeared. Trials were separated by an inter trial interval (ITI) to allow fUSI signal to return to baseline. **b**, Quantitative image similarity metrics for the reduced-time approach. VARS-fUSI performed similarly to reference model, Oracle-fUSI, and outperformed DeepfUS and SVD-fUSI. **c**, Qualitative example images of each models’ performance using the reduced-time approach. **d**, The decoding accuracies over time across a trial for all methods using the reduced-time approach. Full DOP information was used from all 10 reconstructed DOP images. **e**, Statistically significant task-correlated activation maps with *p <* 0.05 of each method using the reduced-time approach calculated by performing voxelwise t-tests with Bonferroni multiple comparisons correction. **Note, the ground-truth activation map only displays voxels with a p-value *<* 0.01 to reduce speckling due to all other models having dampened signal. **f**, Quantitative image similarity metrics for the reduced-sampling approach. **g**, Qualitative example images of each models’ performance using the reduced-sampling approach.**h**, The decoding accuracies over time across a trial for all methods using the reduced-sampling approach. **i**, Statistically significant task-correlated activation maps with p*<* 0.05 of each method using the reduced-sampling approach calculated by performing voxelwise t-tests with Bonferroni multiple comparisons correction. **Note, the ground-truth activation map only displays voxels with a p-value *<* 0.01 to reduce speckling due to all other models having dampened signal.

For the reduced-time approach, we fine-tuned VARS-fUSI, Deep-fUS, and SVD-fUSI on 32 DOP images captured from the beginning of the H1 session at the original DOP imaging rate using 300 IQ frames, and test performance on new session H2. The results highlight that while SVD-fUSI is sensitive to the threshold used to remove tissue clutter, VARS-fUSI successfully filters out a majority of tissue clutter without removing useful blood flow information and without needing threshold tuning per example (fig. 5b). VARS-fUSI provided a DOP image that was less noisy than SVD-fUSI (fig. 5c). In comparison, Deep-fUS showed worse performance than VARS-fUSI and failed on numerous examples, demonstrated via the high performance variance, particularly in the reduced-time approach (fig. 5b).

In the reduced-sampling scenario, all methods produced one DOP image per ground truth DOP. Although models were initially fine-tuned at 32 frames, we increased the frame count to 64 to improve image quality and preserve behavioral information. At 32 frames, reconstructed images showed significant noise in task-activation maps across all models, likely due to the lower resolution and longer acquisition time in human fUSI (1.65 s per DOP) compared to monkey fUSI (1 s per DOP). Thus, reducing the sampling rate by 10× to 32 frames did not provide sufficient temporal resolution to differentiate between slow and fast signal components. This increase to 64 frames is supported by VARS-fUSI, but not by Deep-fUS, which does not allow frame rate adjustment. Therefore, we compared VARS-fUSI at 64 frames to Deep-fUS at 32 frames. At 64 frames, VARS-fUSI outperformed Deep-fUS and Oracle-SVD in both performance metrics and visual image quality(fig. 5f,g). As in the reduced-time approach, Deep-fUS performed significantly worse than VARS-fUSI (fig. 5g).

#### Human experiment decoding

Similar to the aforementioned monkey experiments, we demonstrated that VARS-fUSI can capture behavioral information in a human subject using the same analysis used for monkey decoding.

fUSI data was recorded from the primary motor cortex of a human participant with an acoustic cranial window implant while they performed randomized instructed movement tasks in block task format (see Methods for more task information). Movement effectors included the right index finger, right wrist, lip, and tongue (fig. 5a). DOP data was processed and analyzed offline. Two-sample t-tests of voxel-wise activity during the task vs. baseline were used to map significant task-correlated voxels (*p <* 0.05). The same offline decoding pipeline that was used for the above monkey experiments was then used to decode cued movement effectors.

For the reduced-time approach, one image was reconstructed per 32 consecutive IQ frames from the original 300 IQ frames. This resulted in 10× more DOP images compared to the original imaging framerate. For the reduced sampling approach, the methods constructed one image within the same amount of time as the original 300 IQ frames but using 20% of the sampling rate (64 frames).

We then compared the decoding performance and statistical maps of VARS-fUSI with the ground truth, SVD-fUSI, and Deep-fUS. For decoding, we combined data from five different experiment sessions (table S2). We calculated decoding performance over time from one DOP in time across a trial.

In the reduced-time scenario, VARS-fUSI outperformed Deep-fUSI in decoding accuracy, though it performed slightly below ground truth and SVD-fUSI (fig. 5d). This is likely due to the lower signal-to-noise ratio within human fUSI images compared to monkey fUSI images as well as the complex nature of the task. We also observed that VARS-fUSI produced activation maps similar to ground truth (fig. 5e). We note that the signal was dampened by VARS-fUSI and thus did not have as extensive statistically significant regions of activation. Finally, we verified that regions in the VARS-fUSI activation map that did not align with the ground truth still possessed neural activity waveforms similar to the ground truth but were lower in average percent change in activity (fig. S8).

For the reduced-sampling approach, VARS-fUSI had comparable decoding performance to ground truth, SVD-fUSI, and Deep-fUS (fig. 5h), VARS-fUSI had better activation representation than Deep-fUS. Specifically, using the same statistical testing as above, we observed that VARS-fUSI produced activation representations similar to the ground truth and SVD-fUSI, while Deep-fUS’ representations were much noisier (fig. 5i), challenging the interoperability of Deep-fUS’ decoding.

## Discussion

Functional ultrasound imaging (fUSI) is an emerging neurotechnology with strong potential for long-term, epidural neurorecording during active movement and behavior. fUSI offers higher sensitivity and spatial resolution than fMRI and EEG, while being less invasive than microelectrode recordings and providing a broader field of view but with lower temporal resolution^1^. A major limitation of fUSI is its reliance on many compounded frames to construct a single Doppler power (DOP) image, making acquisition computationally inefficient. Thousands of pulses per DOP increase storage and transfer demands and can cause probe heating during extended sessions, especially with more probe elements. These challenges will intensify with advances such as 3D volumetric imaging, which further raise data requirements per image.

We propose variable sampling for fUSI (VARS-fUSI), a deep-learning solution that reduces the number of frames needed to reconstruct high-quality fUSI images. VARS-fUSI achieves 90% frame reduction while preserving DOP image reconstruction quality. We explore two approaches for efficient imaging with fewer frames: a) reducing the acquisition time (reduced-time) and b) lowering the frame sampling rate (reduced-sampling) per DOP image. Designed for spatiotemporal processing via neural operators, VARS-fUSI adapts to changes in in-phase and quadrature (IQ) sampling rates without compromising image quality. While hardware optimization alone is insufficient, existing deep-learning methods fail to incorporate spatiotemporal filtering, adapt to IQ sampling variations, or assess the usability of reconstructed images for behavioral decoding^14^. VARS-fUSI overcomes these limitations, offering state-of-the-art performance across sessions, planes, and species. VARS-fUSI generates high-quality DOP images from significantly fewer frames and preserves decodable behavioral information, as validated by comparable decoding accuracies.

VARS-fUSI reconstructs high-quality DOP images from limited IQ signals, achieving visual quality and similarity metrics comparable to the SVD-based fUSI standard, while requiring far less time and data. This reduces the need for computationally intensive processing and filtering^13^. VARS-fUSI consistently outperforms deep learning methods such as Deep-fUS^14^. We attribute this generalization to its architecture, which leverages neural operators that respect the complex and temporal data structure, unlike Deep-fUS. VARS-fUSI supports flexible data acquisition rates and durations, a key advantage over existing data-driven approaches. This flexibility, enabled by its temporal neural operator design, makes it broadly applicable. Training and fine-tuning can be done efficiently on a single GPU. Overall, VARS-fUSI is a practical and user-friendly tool for producing DOP images from reduced IQ frames, with wide utility across fUSI experiments without requiring substantial modifications to the acquisition pipeline.

Beyond structural image similarity metrics, VARS-fUSI is the first deep-learning method to reconstruct DOP images with flexible framerates from reduced IQ frames while preserving accurate behavioral decoding. While Deep-fUS produces task-evoked functional activation maps similar to SOTA methods, its performance degrades with increasing data sparsity and does not assess its ability to decode complex behaviors, focusing only on task-correlated activity in visual-stimulation tasks. VARS-fUSI achieves high decoding accuracy, comparable to SVD-based ground truth, across various behavioral tasks in NHPs and humans. Unlike other models with inconsistent quality, VARS-fUSI maintains robust performance across datasets, ensuring stable decoding accuracy. The reduced-time approach offers a higher imaging rate, potentially reducing BCI latency. In the reduced-sampling approach, VARS-fUSI enables imaging and behavioral decoding while reducing data acquisition costs by 90%.

VARS-fUSI offers task-correlated activation maps comparable to ground truth through voxel-wise statistical analysis, confirming spatially specific task-related activity. However, we caution against directly using VARS-fUSI for behavioral mapping due to noise, signal dampening, and potential signal hallucination, especially at lower sampling rates. Despite this, VARS-fUSI reliably constructs DOP images from fewer data without significantly compromising behavioral information, demonstrating its potential to enhance real-time fUSI-based BCIs by increasing imaging framerate, reducing input sampling, and lowering computational and memory costs.

It is important to note that VARS-fUSI does not enhance image quality or decodability relative to the standard being currently used but improves the computational efficiency of fUSI acquisition and processing by reducing the collection wait time (allowing for increased imaging framerate) or by reducing the sampling rate. VARS-fUSI outperforms SVD-based fUSI when the same amount of limited data is used. However, compared to the ground-truth DOP, VARS-fUSI offers an approximated reconstruction of the ground-truth DOP image. Lastly, VARS-fUSI is a deep-learning framework; hence, it still requires a training stage and a dataset consisting of IQ/DOP pairs.

To conclude, VARS-fUSI is a SOTA deep learning platform that enables efficient fUSI imaging from limited time frames. It reconstructs high-quality DOP images from fewer IQ frames—using less temporal data or lower sampling rates—allowing comparable image quality with reduced resources. This is a key advancement for fUSI neurotechnology. As volumetric imaging grows, computational and storage demands will become more limiting. The high pulse counts in 3D imaging risk exceeding FDA thermal limits, causing probe heating and storage challenges. VARS-fUSI can help mitigate these issues by reducing pulse count, sampling rate, and data per DOP image. Overall, VARS-fUSI is well suited for any future fUSI setting where computational efficiency is critical, e.g., fUSI-BCI systems for restoring mobility, therefore expanding the reach and utilization of fUSI.

## Methods

### Notations

Scalars, vectors, and matrices/tensors are denoted by non-bold-lower case *a*, bold-lower-case ***a***, and bold-upper-case letters ***A***, respectively. We denoted the conjugate transpose of matrix ***A*** by ***A***^*^. We let *i* = 1, …, *n* index the number of examples (i.e., pairs of IQ frames and DOP image data). We denoted the measured beamformed IQ frames by ***Y*** ^*i*^ ∈ ℂ^*H×W ×T*^, and its corresponding desired doppler image by ***X***^*i*^ ∈ ℝ^*H×W*^.

### Training pairs

Let 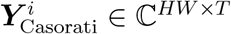 be the 2D space-time Casorati matrix of ***Y***^*i*^. The singular value decomposition (SVD) of 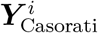 is

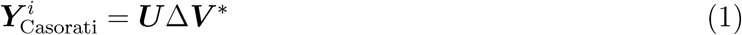

where Δ ∈ ℝ^*HW ×T*^ is a non-square diagonal matrix, containing singular values in descending order of magnitudes. The matrices ***U*** ∈ ℂ^*HW ×HW*^ and ***V*** ∈ ℂ^*T ×T*^ are orthonormal matrices containing the singular vectors. To construct a DOP image, we suppressed the tissue signal concentrated on the first singular values of the signal, i.e.,

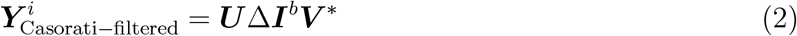

Where ***I***^*b*^ is a block-diagonal matrix with the first *b* diagonal coefficients as 0, and the rest as 1. Denoting the tensor form of the filtered data as 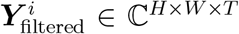, the DOP image is the power average of the decluttered signal as follows:

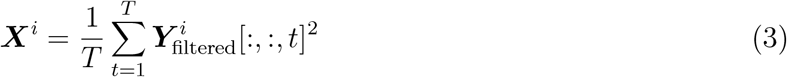

We used this SVD-based approach to produce the pairs 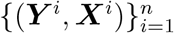 where *T* is set to a large value (e.g., 300 in mice and human, and 250 in monkey experiments). For the reduce-time approach, the training set was constructed based on the above pairs as 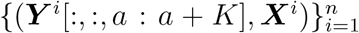, where *a* ∈ [0, *T* − *K*) is an index chosen uniformly at random, and *K* ≪ *T*. For the increased-efficiency approach, the input example was a decimated version of the original input, ***Y*** ^*i*^[:, :, *a* : *b* :], where *a* ∈ [0, *T* − *bK*). We set *b* = 9 and 7 for mice and human experiments with *T* = 300, and monkey with *T* − 250, respectively. The goal was to find a mapping from limited IQ time frames to the desired DOP image ***X***^*i*^.

### Pre-processing

For complex models such as VARS-fUSI, the input data ***Y*** ^*i*^ ∈ ℂ^*H×W ×K*^ and output data ***X***^*i*^ ∈ ℝ^*H×W*^ were standardized as follows

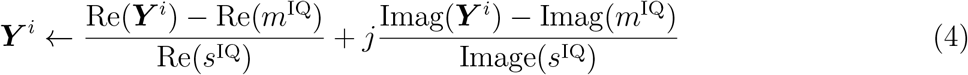

and

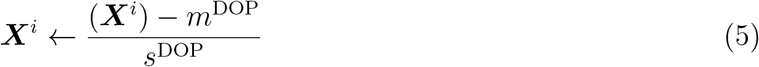

where the standardization constants set {*m*^IQ^, *s*^IQ^, *m*^DOP^, *s*^DOP^} were computed based on the training data as following.

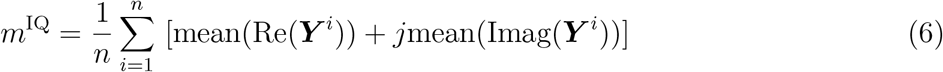

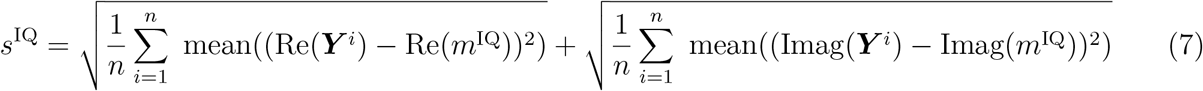

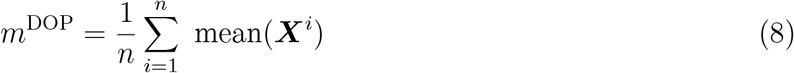

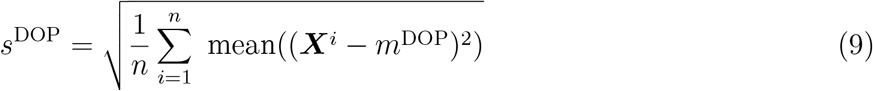

where ‘mean’ stands for the operation to compute one scalar as an average value across the voxels or pixels. Lastly, for models with real values such as DeepfUS, where the model takes only the real part of the complex IQ frames as input, Re(***Y*** ^*i*^), the standardization was

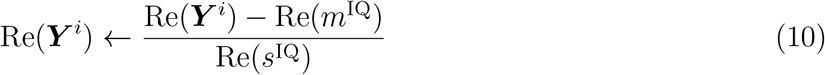

### Augmentation

Prior to standardization, the data pairs were passed through an augmentation pipeline. First, the images/frames were cropped at random to have *H* height and *W* width. Then, three augmentation operations were employed: a) rotations with angle *θ* ∼ Unif([*θ*_1_, *θ*_2_]), b) horizontal flipping of images and c) random vertical flipping of images. Each augmentation occurred at random with a probability of 0.5. We set *H* = *W* = 128, *θ*_1_ = *−* 10, and *θ*_2_ = 10 for training models on MultiMice. The same augmentation was applied to both input and output.

### Loss function

We used a combination of Mean Absolute Error and Mean Square Error to train the network, i.e.,

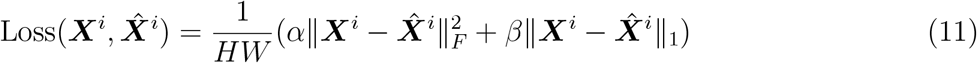

where 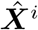 denoted the output of the network. When training was achieved in a batch-based approach, the loss was averaged across the examples within the batch:

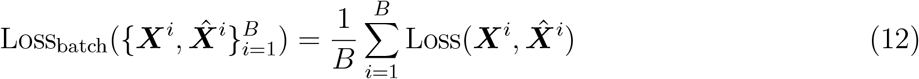

We set *α* = 0.5 and *β* = 0.5 for training models on MultiMice.

### Evaluation metrics

We evaluated the model performances using NMSE (Normalized Mean Square Error), Peak Signal to Noise Ratio (PSNR), and Structural Similarity Index Measure (SSIM), defined below. Both NMSE and SSIM range between 0 and 1.

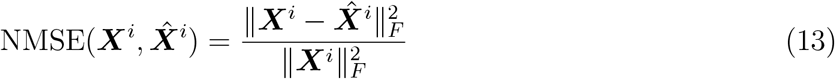

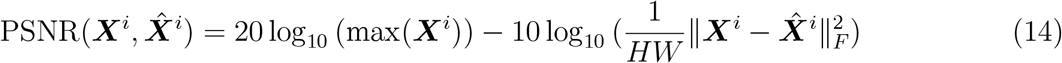

For SSIM 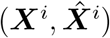, we used the python package `pytorch_msssim` to compute SSIM using a window size of 7. When reporting these numbers on a set, we reported their average across the set. For example, for SSIM, that would be

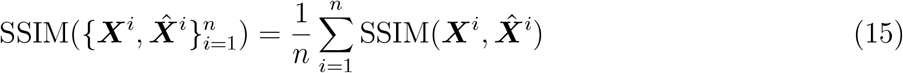

Moreover, prior to computation of PSNR, we normalized the DOP images into a valid dynamic range as follows

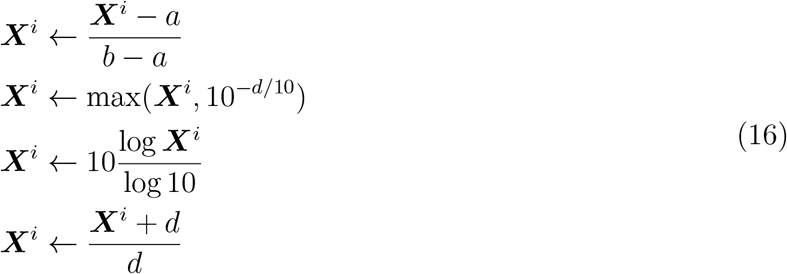

where 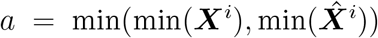 and 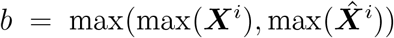. We set *d* = 40 when reporting results. For SSIM, only the first step of normalization was used.

### Training

We trained the models on the MultiMice training set for 600 epochs using Adam optimizer with a learning rate of 10^*−*4^, and weight decay of 10^*−*5^. We used a gradient clip of size 5.0 during training. The batch size was set to 1.

### Fine-tuning

We used a similar procedure as training to fine-tune an already trained model. The only difference was that the model was “warmed-up” from the weights trained on MultiMice and then trained on limited data for a limited time. table S1a describes the details of parameters for finetuning of the models.

### Runtime computation

Runtime is divided into two components: collection-wait-time (cwt), which includes frame acquisition, beamforming, and data storage, and processing-wait-time (pwt), which involves compounding frames and SVD. The cwt is the primary runtime bottleneck compared to the pwt. To obtain the ground truth image, cwt is 2.2 seconds (corresponding to *T* = 300 frames at a 500 Hz IQ sampling rate) and pwt is 0.0949 seconds. Our reduced-time approach enables an approximately 90% reduction in cwt by collecting only *K* = 32 frames at 500 Hz, yielding a cwt of 32*/*300 × 2.2 = 0.23 seconds for all deep learning-based methods discussed. In monkey experiments, reducing IQ frames from 250 to 32 for a single DOP image lowered cwt by 87.2%, effectively addressing latency and hardware efficiency limitations in real-time fUSI for BCIs^7^. The pwt was computed on an NVIDIA GeForce RTX 4090 GPU. SVD computation and related data processing were implemented via a dataloader operating on CPU, and the deep learning frameworks operated on GPU. The reported pwt number is the average runtime of processing 1 DOP image, computed using 1000 DOP in a for-loop; the pwt combines the runtime of the dataloader and the model processing.

We observe that SVD-fUSI using *K* limited IQ frames has lower pwt (0.0055 s) than VARS-fUSI (0.0230 s); however, we note that VARS-fUSI has better image quality metrics than SVD-fUSI. The pwt of other deep learning frameworks are DnCNN (0.0026 s), Deep-fUS-base (0.0031 s), and Deep-fUS (0.0031 s). Although VARS-fUSI pwt is a bit longer than the other deep learning frameworks due to its spatial guidance and delicate design of temporal filtering using Fourier neural operators (FNO)s^16,17^, VARS-fUSI has the best imaging quality among all, and it is still much faster than full-frame *T* SVD with a pwt of 0.0949 s.

Combining the cwt and pwt, the total effective runtime of VARS-fUSI would be 0.2530 s. This runtime is comparable to other deep learning frameworks and significantly faster than the ground-truth method (2.2949 s) performed at full collection wait time with SVD processing. The substantial reduction in cwt (≈ 90%) and pwt (≈75%) results in a remarkable speedup, enabling DOP imaging every 0.2525 s, compared to 2.2949 s for the ground truth.

We also examined the runtime information on VARS-fUS in the absence of spatial guidance. For MultiMice inference tests, VARS-fUS without guidance, VARS-fUSI w/o G, took 0.0225 seconds. Hence, for the total runtime, combining cwt and pwt, VARS-fUSI w/o G ran at 0.2525 seconds per image. For the monkey experiments, the finetuning of VARS-fUSI and VARS-fUSI w/o G took 7.15 and 7.05 minutes, respectively with 3600 gradient steps on NVIDIA GeForce RTX 4090 GPU

### Model architectures

VARS-fUSI is a complex model. The neural blocks are explained below.

#### ComplexConv2d

Given a complex input 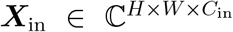, the complex convolution layer, we performed a complex correlation between the input ***x***_in_ and its complex trainable kernels 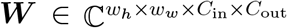 including a bias ***b*** as following

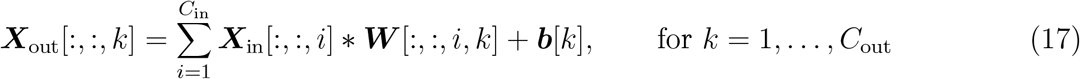

Unless stated, we set *w*_*h*_ = *w*_*w*_ = 3, and padding of size 1 on each side (“same” padding) such that the output had the same *H* × *W* as the input. This layer was built upon torch.nn.Conv2d. The complex computation was achieved as following:

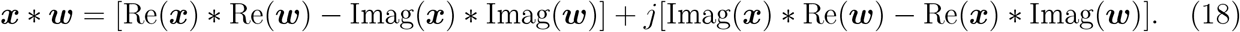

#### ComplexConvTranspose2d

This module is similar to `ComplexConv2d` with the only difference being that it uses convolution with stride of 2 in place of correlation, followed by a ReLU activation function. This was built on torch.nn.ConvTranspose2d.

#### SpatialComplexConv

This block contains the sequence of `ComplexConv2d, ReLU, Dropout2d,` and skip connections, as shown in fig. S1c. This block was designed for spatial processing and is similar in general structure to building blocks in Deep-fUS. This spatial processing runs in parallel for all input time frames and batches. The *i, o* denotes the number of input and output channels, respectively.

#### SpatialConv

This block has a similar architecture to `SpatialComplexConv`, except it does real-valued computations.

#### SpectralTemporalOperator

This block (fig. S1d) takes the input to the frequency domain using fast Fourier transform `(torch.fft.fftn)` across the time dimension, followed by a trainable spectral filtering, removing high-frequency modes. The processed data is then transformed back to the time domain and passed through a `LeakyReLU` activation function. This block also contains a skip connection from the input to the output.

#### VARS-fUSI Layer

This layer, appearing in the encoder of VARS-fUSI architecture, first spatially decimates (by 2) the data with `SpatialMaxPool`. This is followed by `SpatialComplexConv` and `SpectralTemporalOperator` for spatial and temporal filtering, respectively.

#### VARS-fUSI T Layer

This layer, appearing in the decoder of VARS-fUSI architecture, spatially expands (by 2) and interpolates the input data with `ComplexConvTranspose2d`. This is followed by a concatenation of the data with feature maps from the encoder layers. The concatenated data is, then, spatially and temporally processed via `SpatialComplexConv` and `SpectralTemporalOperator`, respectively.

#### MixRealImage

This block fig. S1 uses a Conv3D with kernel sizes of 1 × 1 to linearly combine the real and imaginary components of its complex input.

#### TimeAvg

This block fig. S1 performs a simple averaging across the time frames within its input tensor.

VARS-fUSI architecture (fig. S1) uses the above blocks for Doppler imaging. It takes limited IQ time frames ***Y*** ∈ ℂ^*T ×H×W*^ as input. ***Y*** is passed through a spatial filtering blocks of `SpatialComplexConv, SpatialConv` and a SVD-fUSI block for spatial guidance. Then, they are linearly added (`SpatialConv` outputs 2 times channels more than `SpatialComplexConv` layer to produce both the real and imaginary components required for the complex addition). The signal is then passed through `SpectralTemporalOperator` and then processed by a UNet architecture whose encoder and decoder blocks are modeled by `VARS-fUSI Layer` and `VARS-fUSI T Layer`. The output of the UNet with dimensions (*K* ×*C*×*H* ×*W*) is processed with a ComplexConv2d mapping the *C* channels to 1. This is followed by `MixRealImage` and `TimeAvg` to construct the final DOP image of dimension *H* × *W*.

### Background on neural operators

Neural operators (NOs) are deep learning models that learn a mapping between function spaces^16,17,34^. They are widely being used for finding solutions to partial differential equations (PDEs) widely appearing in science and engineering. Neural operators have found their applications in many disciplines such as seismic signal processing^24^, computational fluid dynamics^22^, weather forecasting^25^, and lithography modeling^35^. The notable property of neural operators is their parameterization can be used at different discretizations. Fourier neural operators (FNOs)^16^ is one of the popular neural operators that parameterize an operator integral kernel in the Fourier domain; hence, they perform global convolution in the original domain.

### Background on UNet

UNet is currently the most popular neural network architecture for parameterization of mappings to be learned from data. It was first introduced in 2015^26^, and since then, its variants have been used in a wide range of applications such as speech enhancement^36^, image generation with diffusion models^37^, and image-restoration tasks when combined with transformers^38^. Two works extended UNet into functional mapping with neural operators and proposed U-shape neural operators for solving PDEs^27,39^.

### Background on singular value decomposition

Ultrasound images consist of signal reflections from the tissue and the blood. Ultrasound imaging techniques aim to remove the clutter caused by tissue from the blood flow. SVD is widely used in this regard^10,11,40^. They compute the SVD of the given data matrix and zeroe out the largest singular values. Given the physics of ultrasound, this filters the tissue signals, containing highly correlated reflections across time frames. Similar techniques have been used for functional ultrasound imaging to filter out the spatially coherent part of the signal corresponding to clutter^5,41^.

### Baseline method of SVD-based filtering

SVD-based fUS is a popular approach for generating DOP images through de-cluttering. The method first computes the SVD of the IQ tensor to exploit spatiotemporal patterns. It then discards the highest contributing components, which contain highly correlated patterns consistent along the time axis. This step removes clutter caused by tissue motion, which is concentrated on first highest singular vectors. The filtered data is subsequently power-averaged to construct the DOP image. We used this approach to construct images for both ground-truth, which used *T* frames for decluttering and *T* frames to compute the DOP, and SVD-fUSI, which performed decluttering and DOP imaging using a lower number of frames, *K*. Lastly, we used an oracle metric, Oracle-fUSI, which decluttered using the future frames of *T* but computed the DOP using only the first *K* frames. It is important to note that this last method was defined solely for performance comparison and is not practical; if one had access to all *T* frames for decluttering, they would ideally use them all for power averaging. Following the mathematical formulation in the Methods section, these methods are defined in math as following. Given 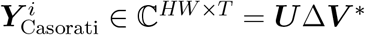, and the filtered signal 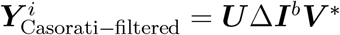, we have

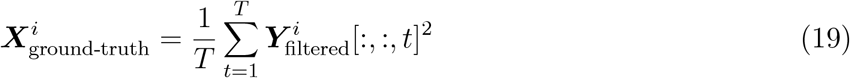

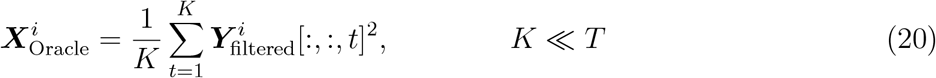

Finally, given 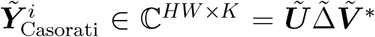 with *K* ≪ *T*, and the filtered signal 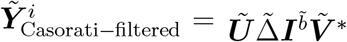, we have

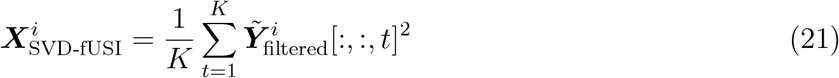

### Baseline method of DnCNN

DnCNN^28^ is a neural network based of residual learning that was originally proposed for image denoising. Recently, DnCNN has been used as learned priors in plug-and-play frameworks and regularization by denoising to solve general inverse problems^42^. The DnCNN that we trained had 17 layers and took only the real portion of the IQ times frames as input. Hence, the incoming and output channels were 32 and 1, respectively. Moreover, the number of features was set to 64 along with a kernel size of 3, a padding of 1, and no bias.

### Baseline method of Deep-fUS

Deep-fUS^14^ is a deep learning framework proposed for functional ultrasound imaging, utilizing a variant of UNet architecture with a focus on input data compression. Deep-fUS discards the imaginary component of the complex data and performs spatial subsampling while reducing the number of time frames, typically ranging from 25 to 125. In its neural network design, Deep-fUS treats time frames as channels, which means it lacks temporal processing capabilities and cannot handle data with a varying number of time frames. In our analysis, we configured Deep-fUS to input a similar number of frames as VARS-fUSI. The training setup and data used for DeepFUS in this paper were identical to those used for VARS-fUSI. We trained two versions of Deep-fUS, one that we refer to as Deep-fUS-base (i.e., UNet in ^14^) and another one as Deep-fUS (i.e., 3D-Res-UNet ^14^). See table S1b for detailed info on the parameters used for Deep-fUS architectures.

### Baseline method of Robust PCA

We attempted a robust PCA-based fUSI approach^43^, as suggested in Solomon et al. 2019^44^, but despite tuning hyperparameters, it performed poorly and required several minutes per image. We attribute this to the reduced number of frames, *K* = 32, compared to the full number of *T* = 300.

### Movement-imitated augmentations for robust BCI decoding

To account for monkey movement during the experiment and increase the robustness of the decoding algorithm to such movements, we incorporated a movement-imitation augmentation step into the BCI decoding pipeline. This simple approach can replace the computationally expensive motion correction algorithms used during offline processing of the experiment session. We introduced an additional data augmentation prior to the other pre-processing steps. Specifically, two rotated and two translated versions of the original Doppler images were concatenated to the original image. The augmentations were randomly selected to fall within a small and reasonable range. For the reported results in fig. S7a, the augmentations used −2 and 2 degrees of rotation and 0.6699*/*0.9656 and −0.7431*/* − 0.9271 x/y translations.

## Data

### Functional ultrasound sequence and recording

Functional ultrasound images rely on ultra-fast Doppler plane waves to detect the backscattered echoes from red blood cells to measure changes in cerebral blood volume. Through neurovascular coupling, we can get a proxy of neural signal.

For recording in mice and non-human primates (NHPs), the ultrasound transducer (Verasonics 128-element miniaturized linear array probe, 15.65-MHz center frequency, 0.1-mm pitch, Vermon, France) was placed on the dura with ultrasound coupling gel. The transducer was consistently placed over the posterior parietal cortex (PPC) using a slotted chamber plug in NHPs. The field of view was 12.8 by 13-20 mm (WxH), which allowed for whole-brain imaging in mice and imaging of multiple cortical regions in NHP including the lateral intraparietal sulcus (LIP, a cortical area that plays a critical role in planning eye movements ^31–33^). Our procedure for fUSI in NHP was identical to that described in Griggs et al., 2024 ^7^: we used a programmable high-framerate ultrasound scanner (Vantage 256; Verasonics, Kirkland, WA) to drive the ultrasound transducer and collect pulse echo radio frequency data. We used a custom plane-wave imaging sequence to acquire the 1 Hz Power Doppler images. We used a pulse repetition frequency of 7500 Hz with 5 evenly spaced tilted angles (−6° to 6°) with 3 accumulations to create one high-contrast compounded ultrasound image. We acquired the high-contrast compound images at 500 Hz and saved the images for offline construction of Power Doppler images. The in-plane resolution was approximately 100 *µm* × 100*µm* with a plane thickness of 400 *µm*.

For recording in our human participant, the ultrasound transducer (Verasonics 128-element miniaturized linear array probe, 7.5 MHz central frequency, 0.3-mm pitch) was placed on top of the skin above the location of the acoustic cranial window. 300 IQ frames were acquired at a 400 Hz framerate and compounded to create one high quality fUSI Doppler image every 1.65 s. Each compounded IQ frame was generated using a pulse repetition frequency of 4000 Hz with 5 evenly spaced tilted angles (−6° to 6°) with 2 accumulations. The transducer was placed over the primary motor cortex (M1) using physical coordinates indicated on an augmented headset. The probe was roughly aligned to MRI using the physical coordinates and then verified functionally through functional mapping of basic motor tasks such as finger tapping. The field of view was 38.4 mm by 49.3 mm (WxH), which allowed for imaging of a wide view of M1. Overall, fUSI acquisition and analysis procedures follow those described in Rabut et al., 2024 ^8^

### Mice subjects

A total of five c57bl/6 mice (6 to 8 weeks old) were imaged. Images were acquired through intact skull and skin after hair removal using a commercial depilatory cream (Nair, Church & Dwigth) without any contrast agent injection. During handling (shaving, positioning) mice were anesthetized with isoflurane (2%) administered in a mixture of 30% O2 and 70% N2. During the resting-state experiment, mice were sedated using dexmedetomidine (Tocris Bioscience). A bolus of 0.10 mg kg-1 was injected subcutaneously, and isoflurane was discontinued after 5 min. Mice were head fixed on a stereotaxic frame to minimize brain motion during imaging.

### NHP subjects

All surgical and animal care procedures were approved by the California Institute of Technology Institutional Animal Care and Use Committee and complied with the Public Health Service Policy on the Humane Care and Use of Laboratory Animals. We implanted one healthy 14-year-old rhesus macaque monkey (Macaca mulatta) weighing 14*kg* with a titanium headpost and placed 24 × 24 mm recording chamber over a craniectomy centered above the left intraparietal sulcus.

### NHP task description

The monkey sat head-fixed facing an LCD screen 30 cm away. Visual stimuli were presented using custom Python 2.7 software based on PsychoPy. The position of the left eye was tracked with an infrared eye tracker at 500 Hz (EyeLink 1000, Ottawa, Canada). Eye position and stimulus information were stored simultaneously for offline analysis.

The monkey performed two variations of the memory-guided saccade task described in Griggs et al., 2024 ^7^ (illustrated in fig. 4a and fig. S6a). At the beginning of the trial, a fixation dot appeared on the screen. To progress the trial, the monkey maintained its gaze fixation on the dot for 5s, after which a peripheral cue was flashed for 500ms. The monkey then continued to maintain center fixation for a 5s memory period until the fixation dot disappeared, at which point he made a saccade to the remembered cue location to obtain a juice reward. The trial was aborted if the monkey broke fixation before the movement period. We used variable durations sampled from a uniform distribution for each task state so the monkey could not anticipate state transitions.

We decoded behavioral parameters (saccade direction and amplitude, fixation location) on a single trial basis by first aligning the fUSI data with the behavioral data so that each fUSI frame was labeled with its corresponding task state and behavioral parameters. The entire image was used for decoding, so each pixel at a given timepoint was a single feature. We decoded the parameters at each time point using a sliding time window comprising the most recent 4 seconds of fUSI data. For example, decoding at the timepoint 5 seconds after the cue used the fUSI images acquired at 2, 3, 4, and 5 seconds, so the input to our decoder had *N* * 4 features where N is the number of pixels in a single Power Doppler image. When decoding, we split the data into train and test folds according to a 15-fold cross-validation scheme. For each fold of data, we scaled the train and test splits by applying a z-score operation fit to the train data, then trained the linear decoder by first applying principal component analysis (PCA) for dimensionality reduction to the train data then applying linear discriminant analysis (LDA) to the results for class separation, keeping 95 percent of the variance from the PCA. We repeated this across all folds and computed the mean decoding accuracy across all folds. We decoded each behavioral parameter (horizontal saccade direction, vertical saccade direction, fixation position, amplitude) separately but always balanced classes across all parameters.

We obtained masks of statistical significance by performing two-sample t-tests of the mean voxel intensity during the memory period for trials from two different conditions (e.g. leftward vs rightward saccade). We report the statistically significant voxels as a mask overlaid on a full power doppler image in fig. 4. For detailed information on animal preparation and implant procedures, the behavioral setup and task, functional ultrasound and recording, and decoding analysis, please see^6,7^.

### Human subject

Our participant was involved in a traumatic brain injury and as a result received a hemicraniotomy to reduce intracranial pressure. After a year, his skull was reconstructed and replaced with a polymeric acoustic skull window (made of polymethyl methacrylate) from Longeviti. The skull replacement is 4 mm thick and lies over a large portion of the left hemisphere, with a 2 mm thinned region of the implant lying over the central gyrus, providing access to M1 and parts of the parietal cortex.

### Human task description

We used fUSI to record changes in CBV from the left hemisphere of M1 while the participant performed instructed delay motor tasks using various parts of the body or “motor effectors.” Motor effectors included the right wrist, right finger, lip, and tongue. During the task, the participant was asked to fixate on a central fixation cross while at rest (16.5 s) and then perform the motion provided at cue onset until the cue disappeared (16.5-24.75 s). The participant was told to repeat the cued motion until the cue disappeared to facilitate a stronger fUSI response. There was then an 8.25 s intertrial interval (ITI) to allow fUSI activity to return to baseline prior to the next trial. To account for the gradual return to baseline in fUSI, the start and end time of each trial was shifted forward by 10 seconds during postprocessing to better capture baseline activity.

We decoded the movement effector used on a single trial basis using a similar pipeline to the monkey decoding experiments. However, rather than using the entire image for decoding, we isolated a specific region of interest based on prior generated activation maps. We decoded the parameters at each time point using the most recent DOP image of fUSI data. When decoding, we split the data into train and test folds according to a 10-fold cross-validation scheme.

We obtained masks of statistical significance by performing voxel-wise two-sample t-tests of the mean voxel intensity during task execution vs. baseline. We report the statistically significant voxels as a mask of p-values overlaid on a full power doppler image in fig. 5.

## Supplementary Figures and Tables

**Table S1:** Finetuning and network parameters.

(a) Fine-tuning parameters on experiment for Mouse X, Monkey, and Human.
| Mouse X |  |  |  |
| --- | --- | --- | --- |
| H | 128 | W | 128 |
| Number of finetuning data | 2 from M1 | Training iterations | 1600 |
| T | 300 | ground-truth SVD th | 30 |
| K | 32 | limited SVD th | 75 |
| Monkey |  |  |  |
| H | 136 | W | 136 |
| Number of finetuning data | 120 from S1 | Training iterations | 3600 |
| T | 250 | ground-truth SVD th | 38 |
| K | 32 | limited SVD th (reduce-time) | 75 |
|  |  | limited SVD th (reduced-sampling) | 125 |
| Human |  |  |  |
| H | 208 | W | 128 |
| Number of finetuning data | 32 from H1 | Training iterations | 5120 |
| T | 300 | ground-truth SVD th | 20 |
| K | 32 | limited SVD th (reduced-time) | 60 |
|  |  | limited SVD th (reduced-sampling) | 110 |

(b) Network parameters of VARS-fUSI and Deep-fUS.
| VARS-fUSI |  |  |  |
| --- | --- | --- | --- |
| Dropout probability | 0.2 | Residual connections | True |
| Number of encoding/decoding layers | 5 | Nonlinearity | ReLU |
| Number of channels | 16, 32, 64, 128, 256 | Spectral modes | 9 |
| Deep-fUS |  |  |  |
| Dropout probability | 0.2 | Residual connections | True |
| Number of encoding/decoding layers | 5 + (1 Conv3D) | Nonlinearity | ReLU |
| Number of channels | 32, 64, 128, 256, 512 | Input Conv3D | True |
| Conv3D output channel | 4 | Relative kernel size | 0.2 |
| Deep-fUS-base |  |  |  |
| Dropout probability | 0.2 | Residual connections | False |
| Number of encoding/decoding layers | 5 | Nonlinearity | ReLU |
| Number of channels | 32, 64, 128, 256, 512 | Input Conv3D | False |

**Table S2:** Experimental data.

| MultiMice |  |  |
| --- | --- | --- |
|  | ground-truth SVD th | Number of images |
| Mouse 1AAN | 40 | 1,219 |
| Mouse 1ABN | 50 | 539 |
| Mouse 2AAR | 60 | 557 |
| Mouse A | 60 | 38 |
| Mouse B | 60 | 293 |
| Test set | - | 366 |
| New Mouse (Mouse X) |  |  |
|  | ground-truth SVD th | Number of images |
| M1 | 30 | 520 |
| M2 | 30 | 80 |
| Monkey |  |  |
|  | ground-truth SVD th | Number of images |
| S1 | 38 | 10,800 |
| S2 | 38 | 10,800 |
| S3 | 38 | 7,200 |
| Human |  |  |
|  | ground-truth SVD th | Number of images |
| H1 | 20 | 320 |
| H2 | 20 | 310 |
| H3 | 20 | 310 |
| H4 | 20 | 310 |
| H5 | 20 | 310 |

**Figure S1:**
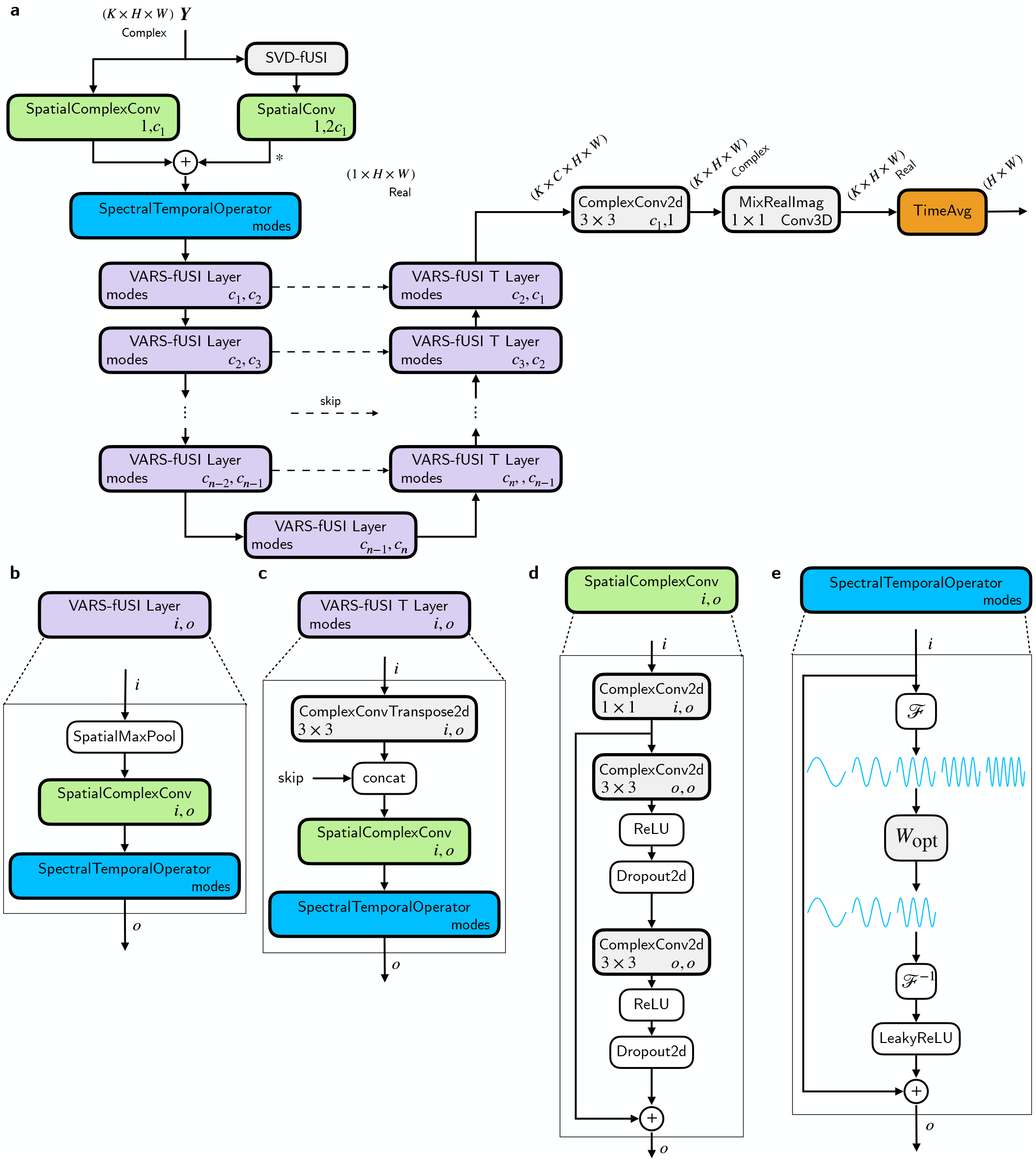
VARS-fUSI architecture and blocks. **a**, VARS-fUSI takes IQ complex frames ***Y***. The data is passed through a decomposed spatial and temporal filtering. It contains a SVD-fUSI step for spatial guidance and a UNet architecture with customized VARS-fUSI Layer and VARS-fUSI T Layer in its encoder and decoder. **b**, VARS-fUSI Layer block. **c**, VARS-fUSI T Layer block. **d**, SpatialComplexConv block. **e**, SpectralTemporalOperator block.

**Figure S2:**
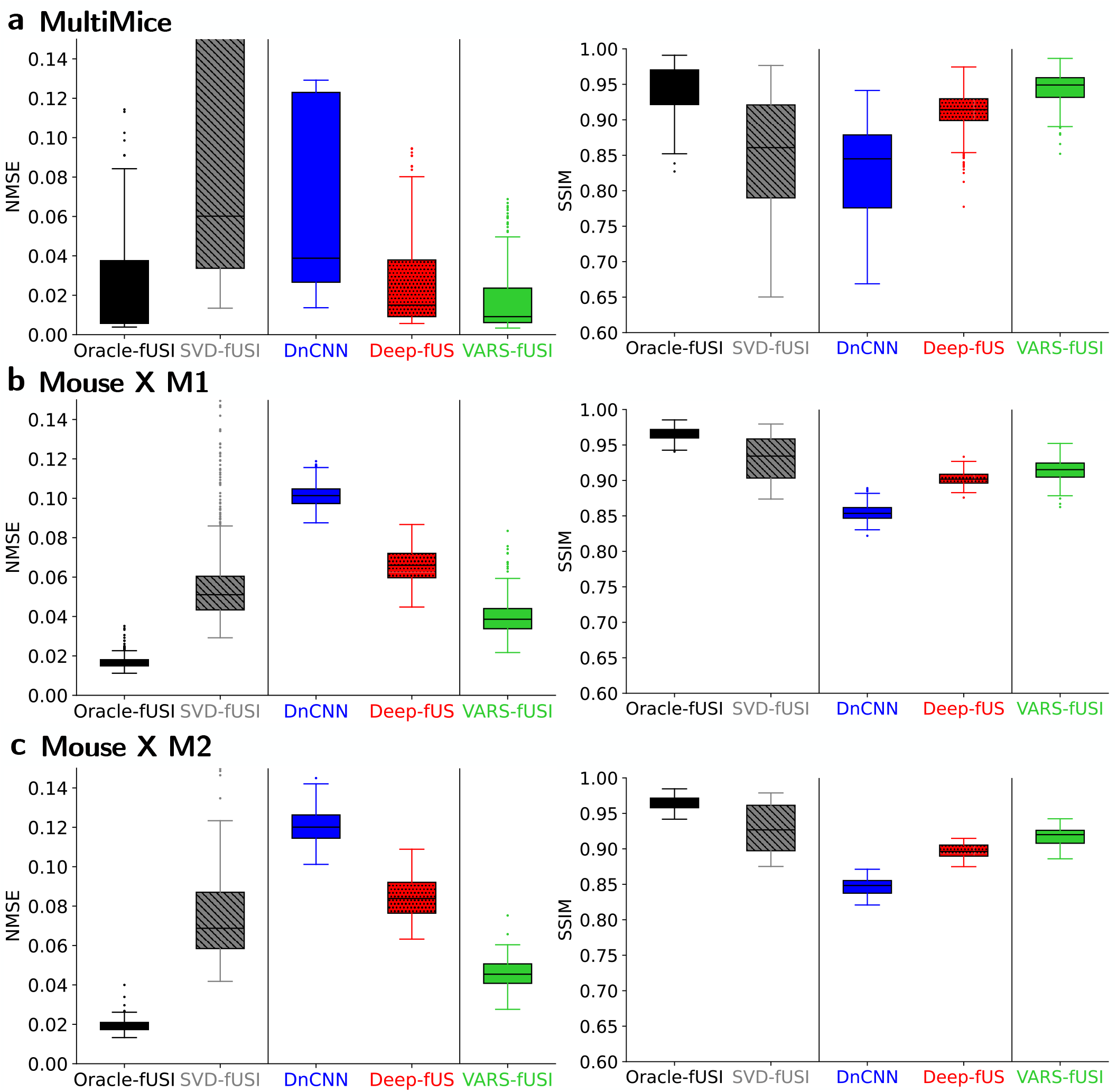
Additional quantification for the outperformance generalization of VAR-fUSI on unseen frames for reduced-time approach. NMSE and SSIM are shown. **a**, MultiMice test set. **b-c**, New mouse Mouse X sessions M1 and M2, respectively.

**Figure S3:**
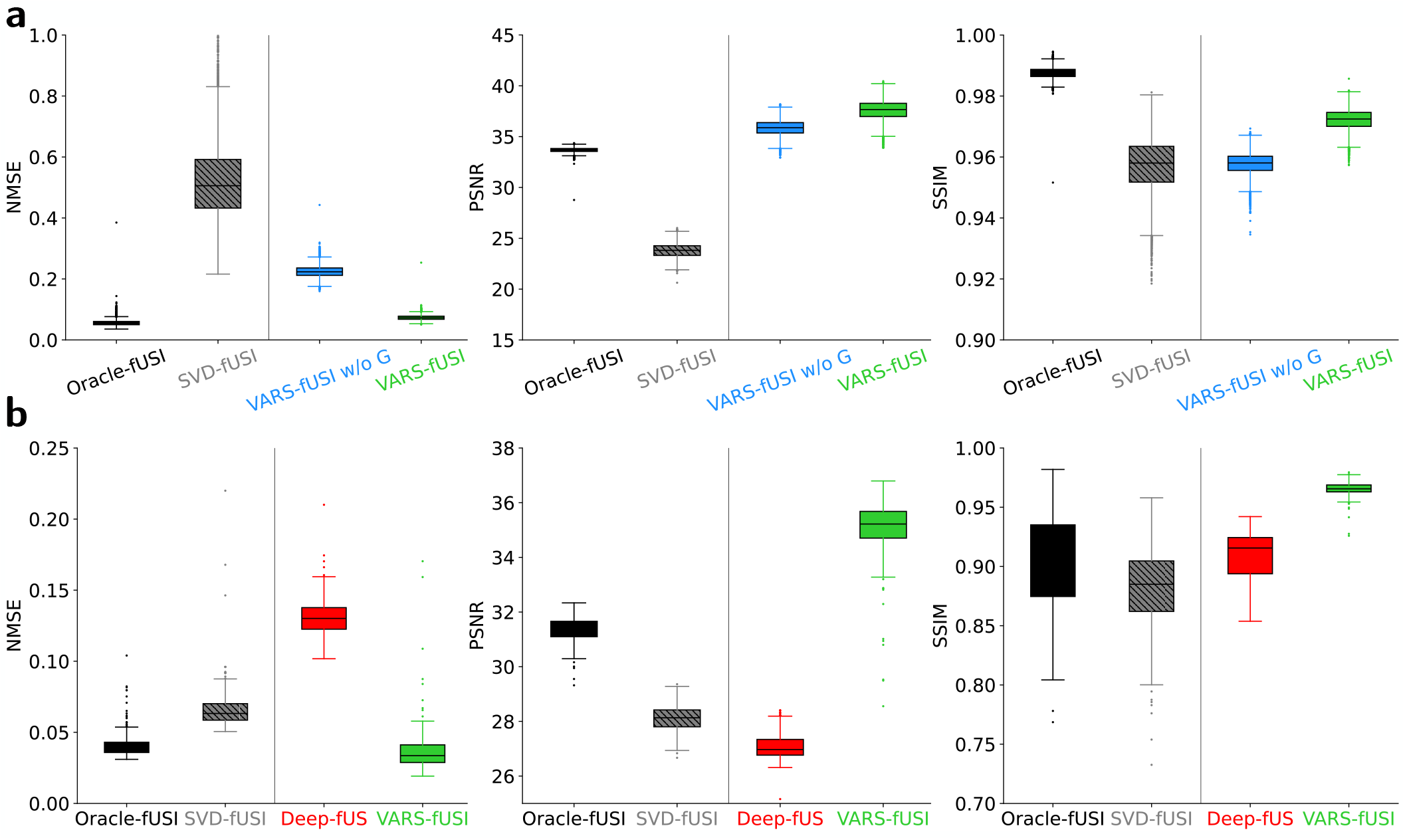
Demonstration of the impact of spatial guidance and limited-regime generalization of VARS-fUSI. **a**, Quantitative performance on the monkey session S2 for reduced-time approach. The figures compare VARS-fUSI with VARS-fUSI. The models are fine-tuned on first 120 images from session S1. **b**, Quantitative performance on human for H2. The figures compare Deep-fUS scratch with VARS-fUSI scratch where “scratch” means the models are not trained on MultiMice set as other results; the models are trained from scratch on only the first 32 DOP images captured during session H1 from Human experiment. The models aimed to produce DOP images from only 32 input IQ frames. From left to right, NMSE, PSNR, and SSIM quantification analysis on session H2 from human experiment. The result highlights the superior generalization of VARS-fUSI compared to Deep-fUS in data-regime.

**Figure S4:**
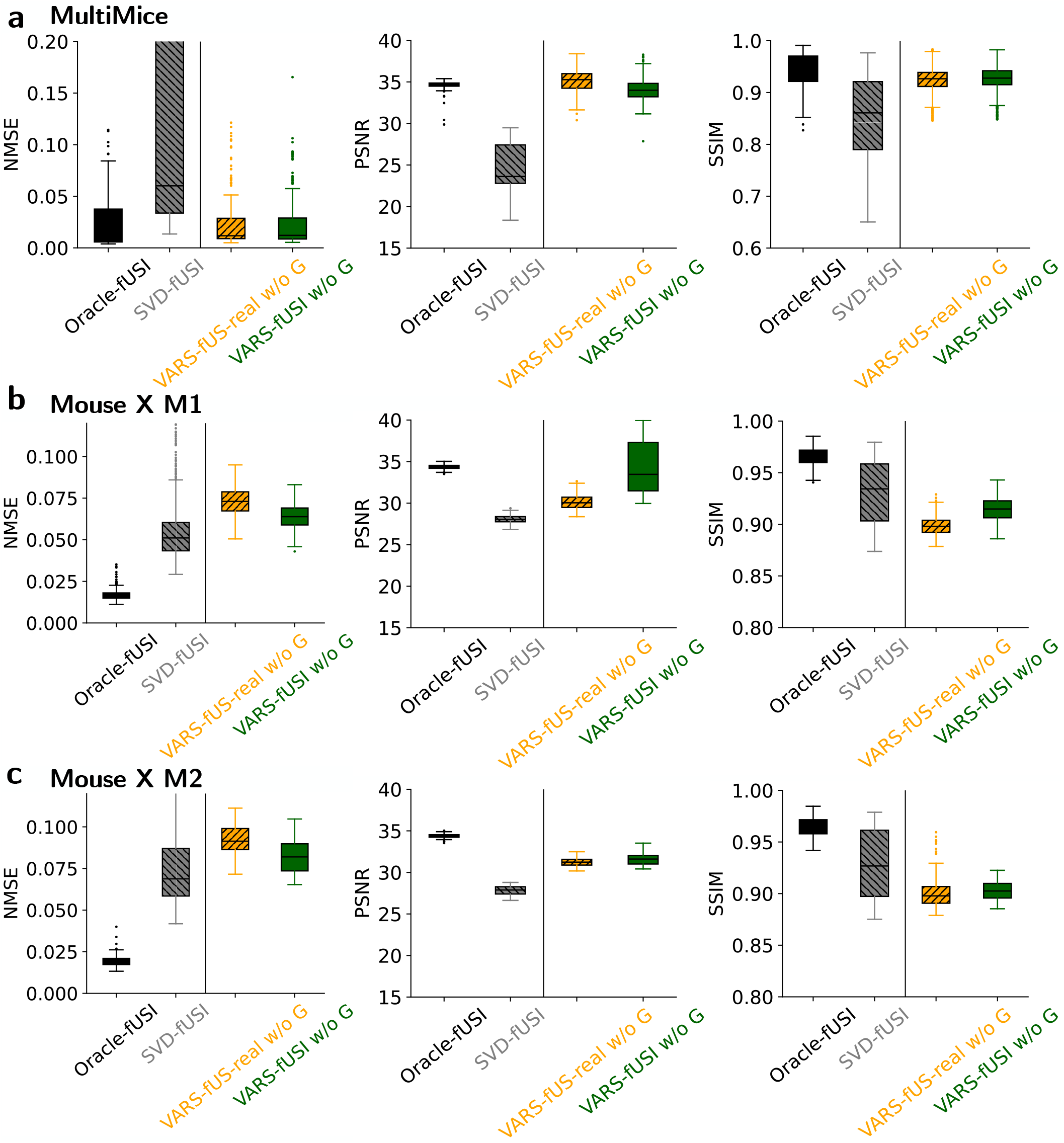
Demonstration of better generalization of complex VARS-fUSI compared to real VARS-fUSI. Frameworks are trained on MultiMice. **a**, Quantitative performance of VARS-fUSI-real w/o G and VARS-fUSI w/o G on Test set of MultiMice in the absence of spatial guidance (w/o G). **b-c**, Generalization performance on the new mouse (Mouse X) for M1 and M2.

**Figure S5:**
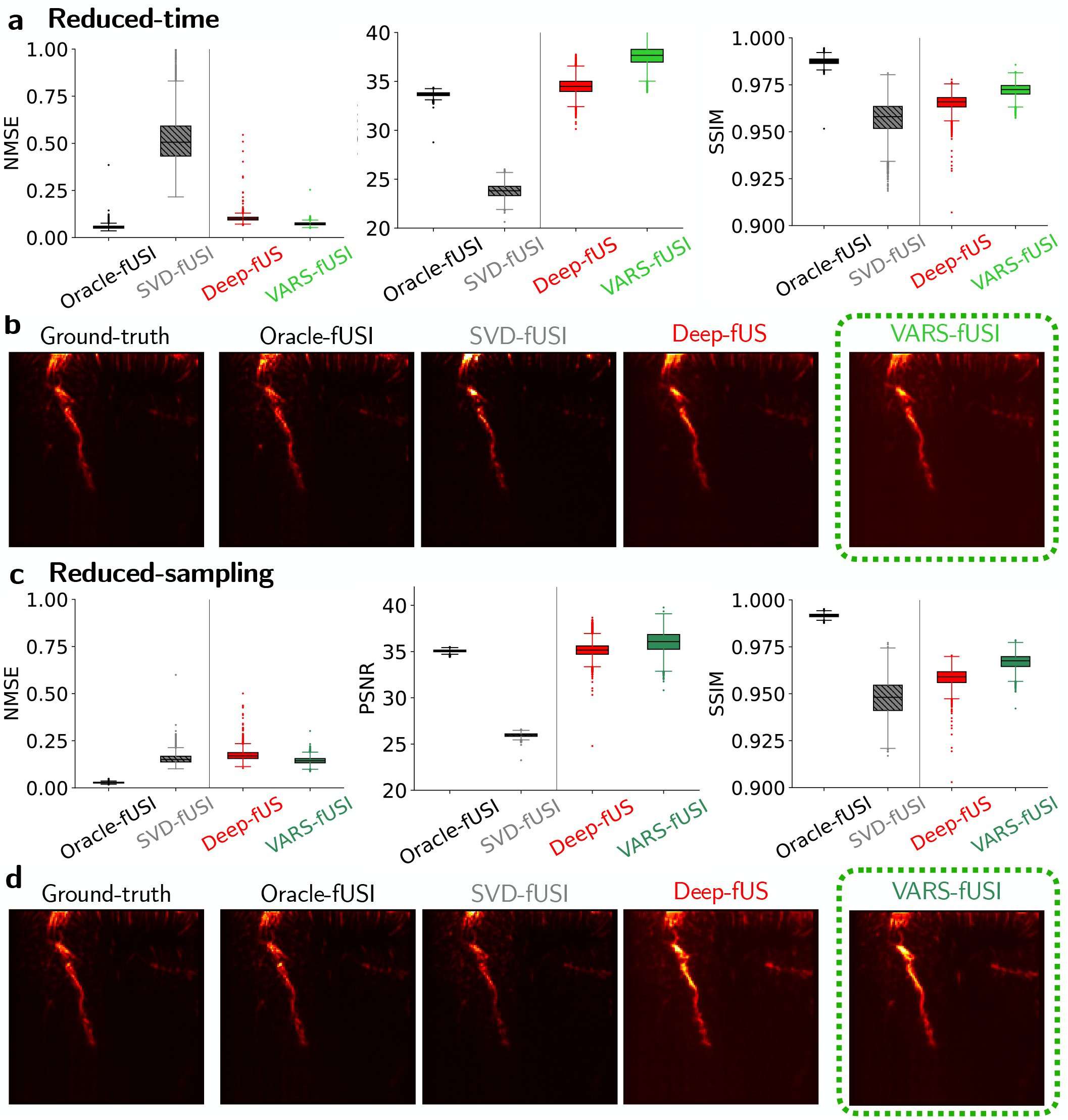
Quantification of imaging on unseen session S2. The models are fine-tuned, on the first 120 DOP images from S1, to create one DOP image using 32 frames as opposed to densely sampled 250 IQ frames. **a-b**, Reduced-time approach. **a**, NMSE, PSNR, and SSIM are shown from left to right for S2. **b**, DOP image example from S2. **c-d**, Reduced-sampling approach. The wait time is similar to the ground-truth, but the data acquisition sampling rate is reduced by a factor of 9*×*. **c**, NMSE, PSNR, and SSIM are shown from left to right. **d**, DOP image example from S2.

**Figure S6:**
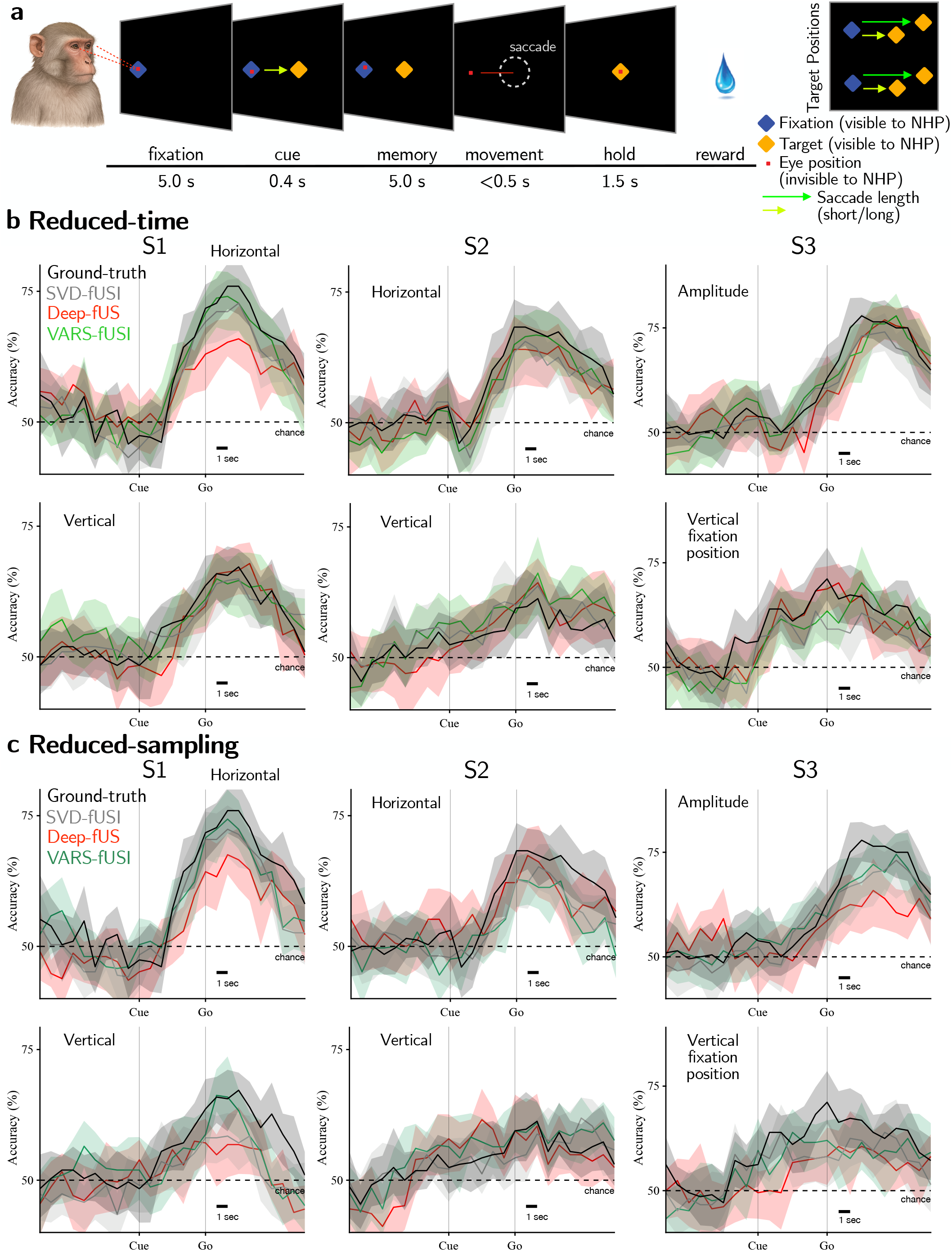
Additional experiments on decoding thoughts from monkey in S1, S2, and S3 sessions. **a**, The experiment setup for S3. **b**, Decoding using reduced-time approach. From left to right: S1, S2, and S3. S1 and S2 shows horizontal (top) and vertical (bottom) decoding. S3 shows amplitude (top) and vertical fixation position (bottom) decoding. **c**, Decoding from images acquired using the reduced-sampling approach. The panels are similar to (**b**).

**Figure S7:**
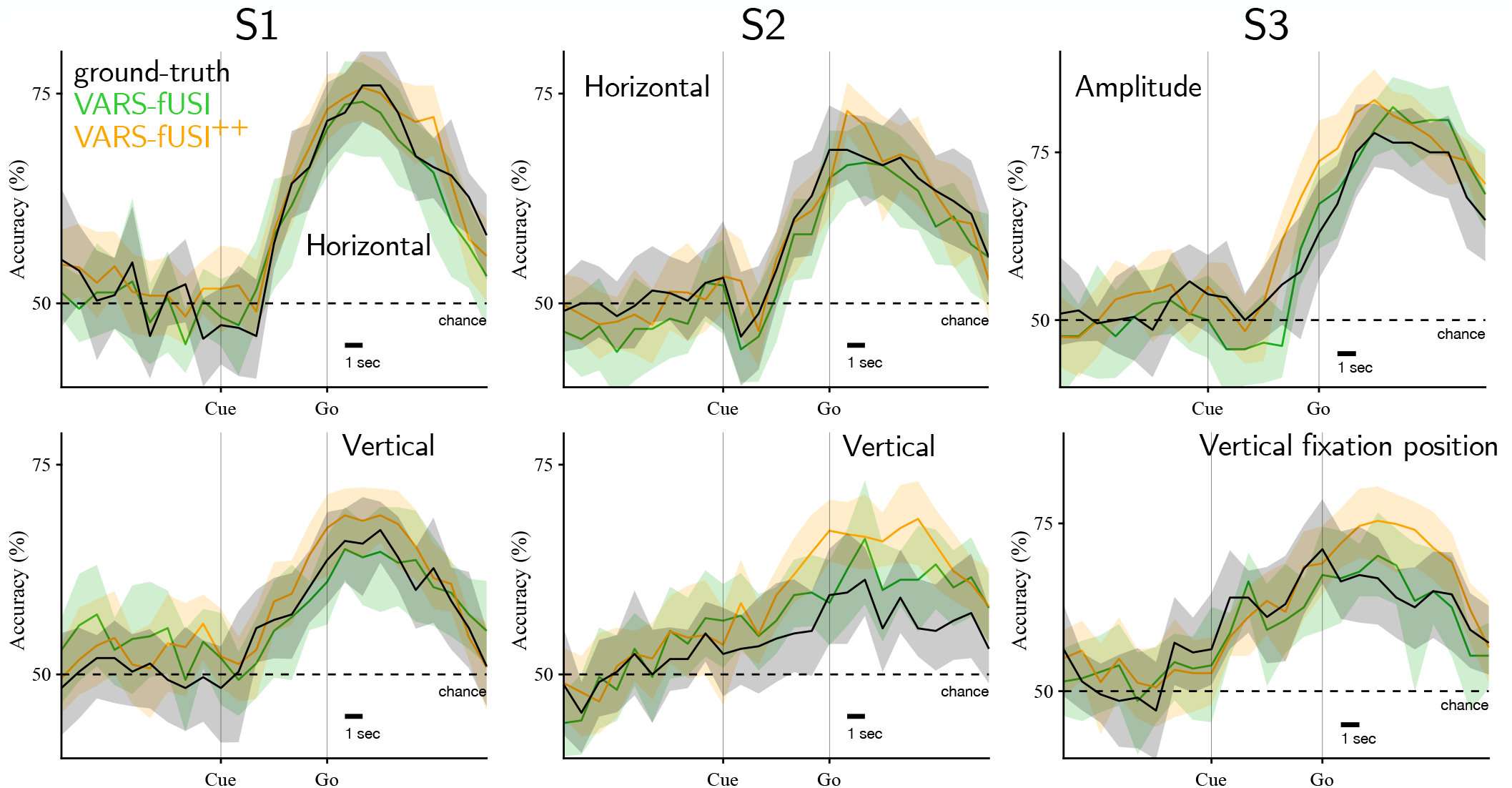
Additional results for monkey decoding. **a**, Motion-imitated augmentations to improve the robustness of decoding thoughts from the monkey for sessions S1, S2, and S3 for the reduced-time approach. VARS-fUSI^++^ stands for VARS-fUSI with the augmentation technique.

**Figure S8:**
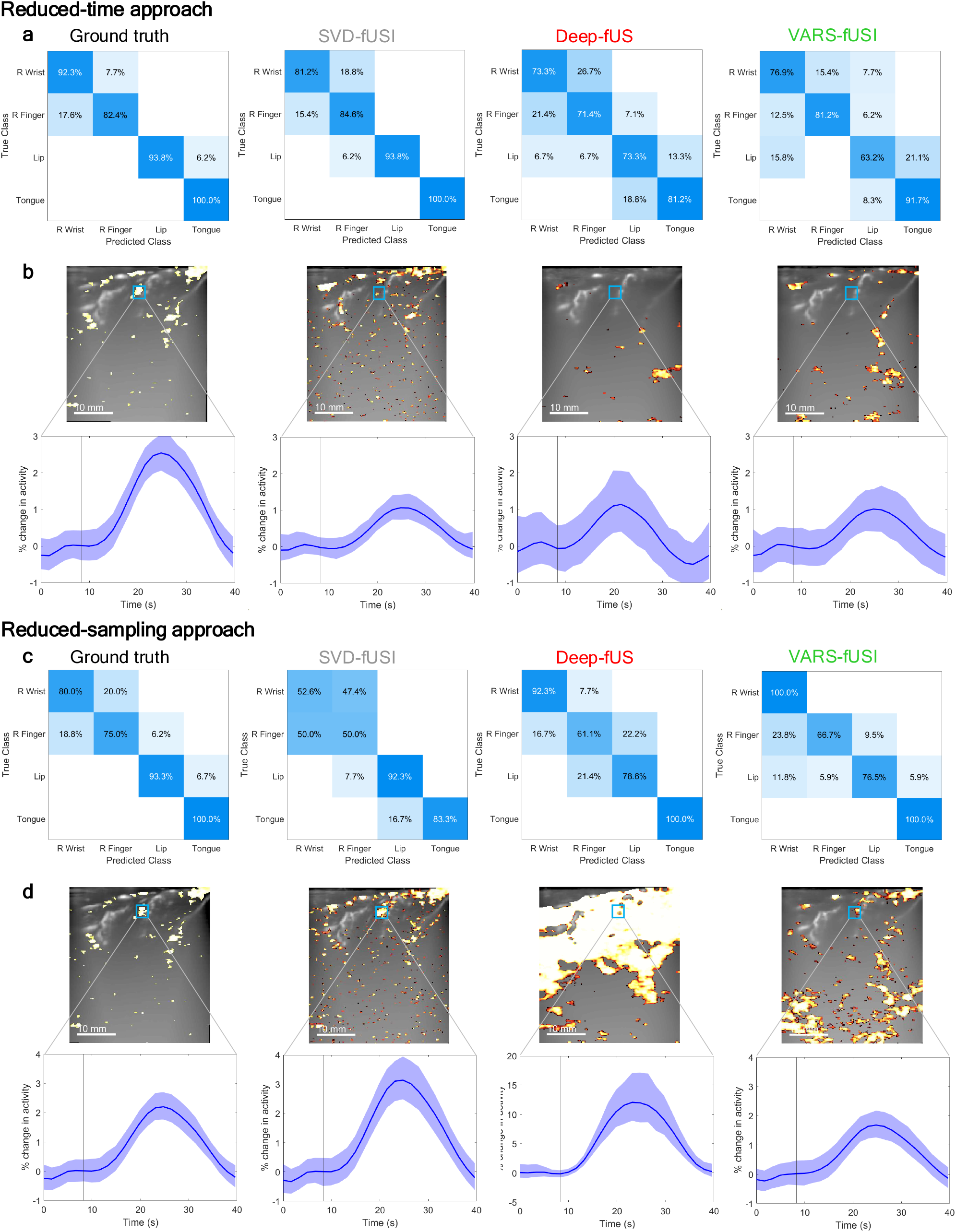
Additional results for human decoding. **a**, Confusion matrices of the best decoding accuracy for each model for the reduced-time approach. **b**, Comparison of fUSI signal waveforms across models in regions of the image that are statistically significant in the ground truth activation map but not for other models using the reduced-time approach. This demonstrates that VARS-fUSI still reconstructs signal appropriately but dampens the signal, reducing the number of voxels that are statistically significant. SVD-fUSI, Deep-fUS, and VARS-fUSI all show dampened signal compared to ground truth. Deep-fUS also shows increased variability in activity across trials **c**, Confusion matrices of the best decoding accuracy for each model for the reduced-sampling approach. **d**, Comparison of fUSI signal waveforms across models in regions of the image that are statistically significant in the ground truth activation map but not for other models using the reduced-sampling approach. VARS-fUSI, similar to the reduced-time approach, is able to reconstruct signal in a similar waveform to ground truth. In comparison, SVD-fUSI and Deep-fUS show increased fUS signal, possibly due to decreased sampling making complete removal of tissue clutter signal difficult.

## Footnotes

1 https://github.com/neuraloperator/varsfusi

